# precisionFDA Truth Challenge V2: Calling variants from short- and long-reads in difficult-to-map regions

**DOI:** 10.1101/2020.11.13.380741

**Authors:** Nathan D. Olson, Justin Wagner, Jennifer McDaniel, Sarah H. Stephens, Samuel T. Westreich, Anish G. Prasanna, Elaine Johanson, Emily Boja, Ezekiel J. Maier, Omar Serang, David Jáspez, José M. Lorenzo-Salazar, Adrián Muñoz-Barrera, Luis A. Rubio-Rodríguez, Carlos Flores, Konstantinos Kyriakidis, Andigoni Malousi, Kishwar Shafin, Trevor Pesout, Miten Jain, Benedict Paten, Pi-Chuan Chang, Alexey Kolesnikov, Maria Nattestad, Gunjan Baid, Sidharth Goel, Howard Yang, Andrew Carroll, Robert Eveleigh, Mathieu Bourgey, Guillaume Bourque, Gen Li, MA ChouXian, LinQi Tang, DU YuanPing, ShaoWei Zhang, Jordi Morata, Raúl Tonda, Genís Parra, Jean-Rémi Trotta, Christian Brueffer, Sinem Demirkaya-Budak, Duygu Kabakci-Zorlu, Deniz Turgut, Özem Kalay, Gungor Budak, Kübra Narcı, Elif Arslan, Richard Brown, Ivan J Johnson, Alexey Dolgoborodov, Vladimir Semenyuk, Amit Jain, H. Serhat Tetikol, Varun Jain, Mike Ruehle, Bryan Lajoie, Cooper Roddey, Severine Catreux, Rami Mehio, Mian Umair Ahsan, Qian Liu, Kai Wang, Sayed Mohammad Ebrahim Sahraeian, Li Tai Fang, Marghoob Mohiyuddin, Calvin Hung, Chirag Jain, Hanying Feng, Zhipan Li, Luoqi Chen, Fritz J. Sedlazeck, Justin M. Zook

## Abstract

The precisionFDA Truth Challenge V2 aimed to assess the state-of-the-art of variant calling in difficult-to-map regions and the Major Histocompatibility Complex (MHC). Starting with FASTQ files, 20 challenge participants applied their variant calling pipelines and submitted 64 variant callsets for one or more sequencing technologies (~35X Illumina, ~35X PacBio HiFi, and ~50X Oxford Nanopore Technologies). Submissions were evaluated following best practices for benchmarking small variants with the new GIAB benchmark sets and genome stratifications. Challenge submissions included a number of innovative methods for all three technologies, with graph-based and machine-learning methods scoring best for short-read and long-read datasets, respectively. New methods out-performed the 2016 Truth Challenge winners, and new machine-learning approaches combining multiple sequencing technologies performed particularly well. Recent developments in sequencing and variant calling have enabled benchmarking variants in challenging genomic regions, paving the way for the identification of previously unknown clinically relevant variants.

## Introduction

PrecisionFDA began in 2015 as a research effort to support FDA’s regulatory standards development in genomics and has since expanded to support all areas of omics. The platform provides access to on-demand high-performance computing instances, a community of experts, a library of publicly available tools, support for custom tool development, a challenge framework, and virtual shared Spaces where FDA scientists and reviewers collaborate with external partners. The precisionFDA challenge framework is one of the platform’s most outward facing features. The framework enables the hosting of biological data challenges in a public facing environment, with available resources for submission testing and validation. precisionFDA challenges, and challenges led by other groups like DREAM (Ewing et al., 2015; Lee et al., 2018; Salcedo et al., 2020, http://dreamchallenges.org) and CAGI (Andreoletti et al., 2019; Hoskins et al., 2017), focus experts around the world on common problems in areas of evolving science such as genomics, proteomics, and artificial intelligence.

The first Genome In A Bottle (GIAB)-precisionFDA Truth Challenge took place in 2016, and asked participants to call small variants from short-reads for two GIAB samples (Zook et al., 2019). Benchmarks for HG001 (a.k.a. NA12878) were previously published, but no benchmarks for HG002 were publicly available at the time. This made it the first blinded germline variant calling challenge, and the public results have been used as a point of comparison for new variant calling methods (Kim et al., 2018). There was no clear evidence of over-tuning methods to HG001, but performance was only assessed on relatively “easy” genomic regions accessible to the short-reads used to form the v3.2 GIAB benchmark sets (Zook et al., 2019).

Since the first challenge, GIAB expanded the benchmarks beyond the easy regions of the genome and improved benchmarking methods. With the advent of accurate small variant calling from long reads using machine learning (Luo et al., 2020; Wenger et al., 2019), GIAB has developed new benchmarks that cover more challenging regions of the genome (Chin et al., 2019; Wagner et al., 2020), including challenging genes that are clinically important (Lincoln et al., 2020). In collaboration with the Global Alliance for Genomics and Health (GA4GH), the GIAB team defined best practices for small variant benchmarking (Krusche et al., 2019). These best practices provide criteria for performing sophisticated variant comparisons that account for variant representations differences along with a standardized set of performance metrics. To improve insight into strengths and weaknesses of methods, for this work we developed new stratifications by genomic context (e.g. low complexity or segmental duplications). The stratified benchmarking results allow users to identify genomic regions where a particular variant calling method performs well and where to focus optimization efforts.

In light of recent advances in genome sequencing, variant calling, and the GIAB benchmark set, we conducted a follow up truth challenge from May to June 2020. The Truth Challenge V2 (https://precision.fda.gov/challenges/10) occurred when the v4.1 benchmark was available for HG002, but only v3.3.2 benchmark was available for HG003 and HG004. In addition to making short-read datasets available (at a lower 35X coverage than the first Truth Challenge), this challenge included long-reads from two technologies to assess performance across a variety of data types. This challenge made use of the robust benchmark tools and stratification BED files developed by the GA4GH Benchmarking Team and GIAB to assess performance in particularly difficult regions like segmental duplications and the Major Histocompatibility Complex (MHC) (Cleary et al., 2015; Krusche et al., 2019; McDaniel et al., 2020). With 64 submissions across the three technologies, the results from this challenge provide a new baseline for performance to inspire ongoing advances in variant calling particularly for challenging genomic regions.

## Results

Participants were tasked with generating variant calls as VCF files using data from one or multiple sequencing technologies for the GIAB Ashkenazi Jewish trio, available through the precisionFDA platform (Fig. 1). Sequencing data were provided as FASTQ files from three technologies: Illumina, Pacific Biosciences (PacBio) HiFi, and Oxford Nanopore Technologies (ONT), for the three human samples. The read length and coverage of the sequencing datasets were selected based on the characteristics of datasets used in practice and manufacturer recommendations (Table 1). Participants used these FASTQ files to generate variant calls against the GRCh38 version of the human reference genome.

**Table 2:** Sequencing dataset characteristics. Read Length - N50 used to summarize PacBio and ONT read lengths. Coverage - median coverage across autosomal chromosomes.

| Technology | GIAB ID | Read Length (bp) | Number of Reads | Coverage |
| --- | --- | --- | --- | --- |
| Illumina | HG002 | 2X151 | 415,086,209 | 35 |
|  | HG003 | 2X151 | 419,192,650 | 35 |
|  | HG004 | 2X151 | 420,312,085 | 35 |
| PacBio HiFi | HG002 | 12,885 | 8,449,287 | 36 |
|  | HG003 | 14,763 | 7,288,357 | 35 |
|  | HG004 | 15,102 | 7,089,316 | 35 |
| ONT | HG002 | 50,380 | 19,328,993 | 47 |
|  | HG003 | 44,617 | 23,954,632 | 85 |
|  | HG004 | 48,060 | 29,319,334 | 85 |

**Figure 1:**
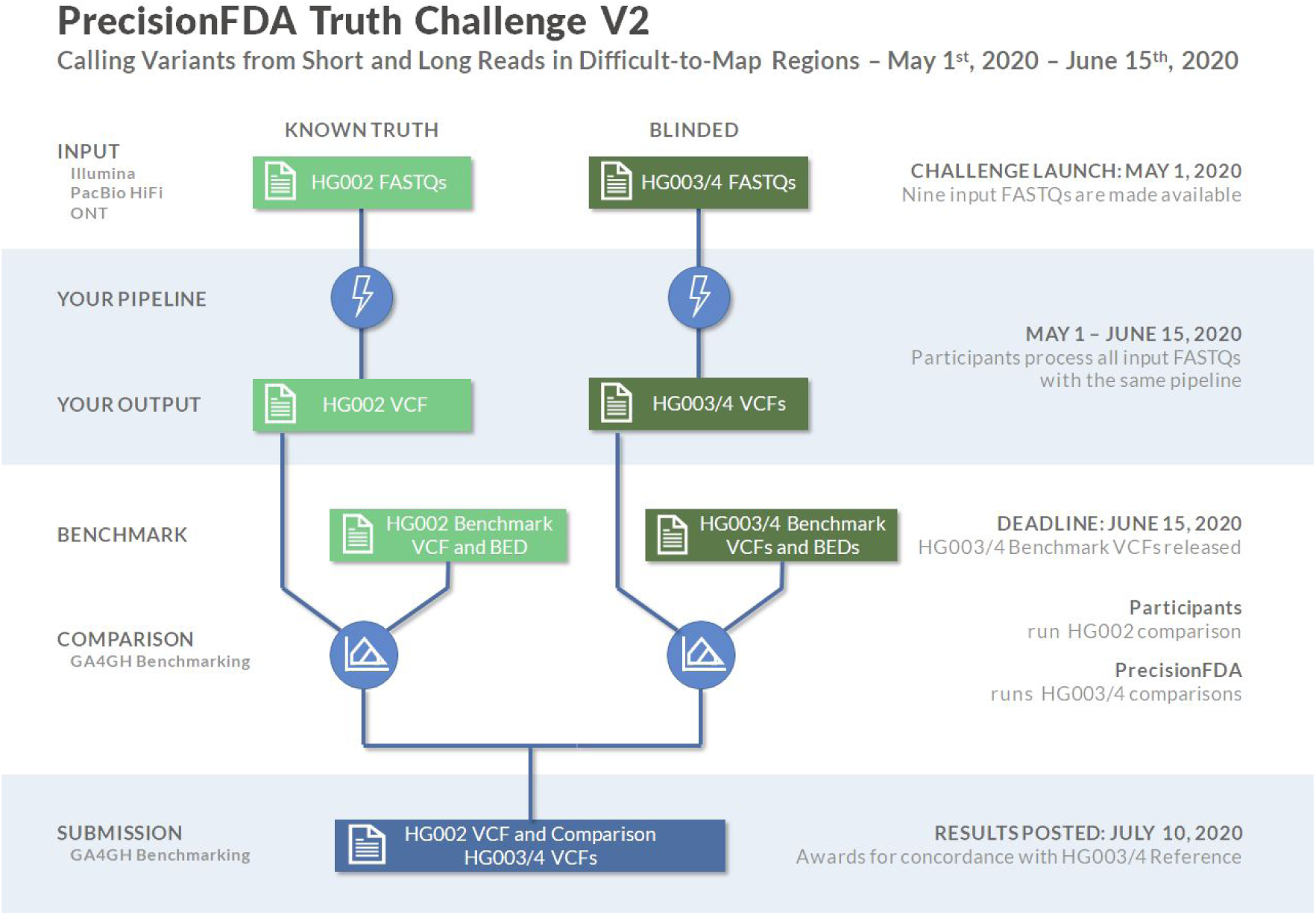
Truth Challenge V2 structure. Participants were provided sequencing reads (FASTQ files) from Illumina, PacBio HiFi, and ONT for the GIAB Ashkenazi trio (HG002, HG003, and HG004). Participants uploaded VCF files for each individual before the end of the challenge, and then the new benchmarks for HG003 and HG004 were made public.

Twenty teams participated in the challenge with a total of 64 submissions, with multiple submissions from a number of teams (Fig.2, Supplemental Table 1). Challenge participants submitted variant callsets that were generated using one or more sequencing technologies, Illumina, PacBio HiFi, and ONT Ultralong (see methods for datasets descriptions). For single technology submissions, Illumina was the most common (24 out of 44), followed by PacBio (17), and ONT (3). PacBio was used in all 20 of the multiple technology submissions, Illumina was used in all but one, and seven submissions used data from all three technologies. Submissions used a variety of variant calling methods based on machine learning (ML; e.g. DeepVariant), graph (e.g., DRAGEN and Seven Bridges), and statistical (e.g. GATK) methods. See supplemental material submission methods for participant variant calling methods. Notably, a majority of submissions used ML-based variant calling methods (Fig. 2B). This was particularly true for long-read and multi-technology submissions, with 37 out of 40 using an ML-based method.

**Figure 2:**
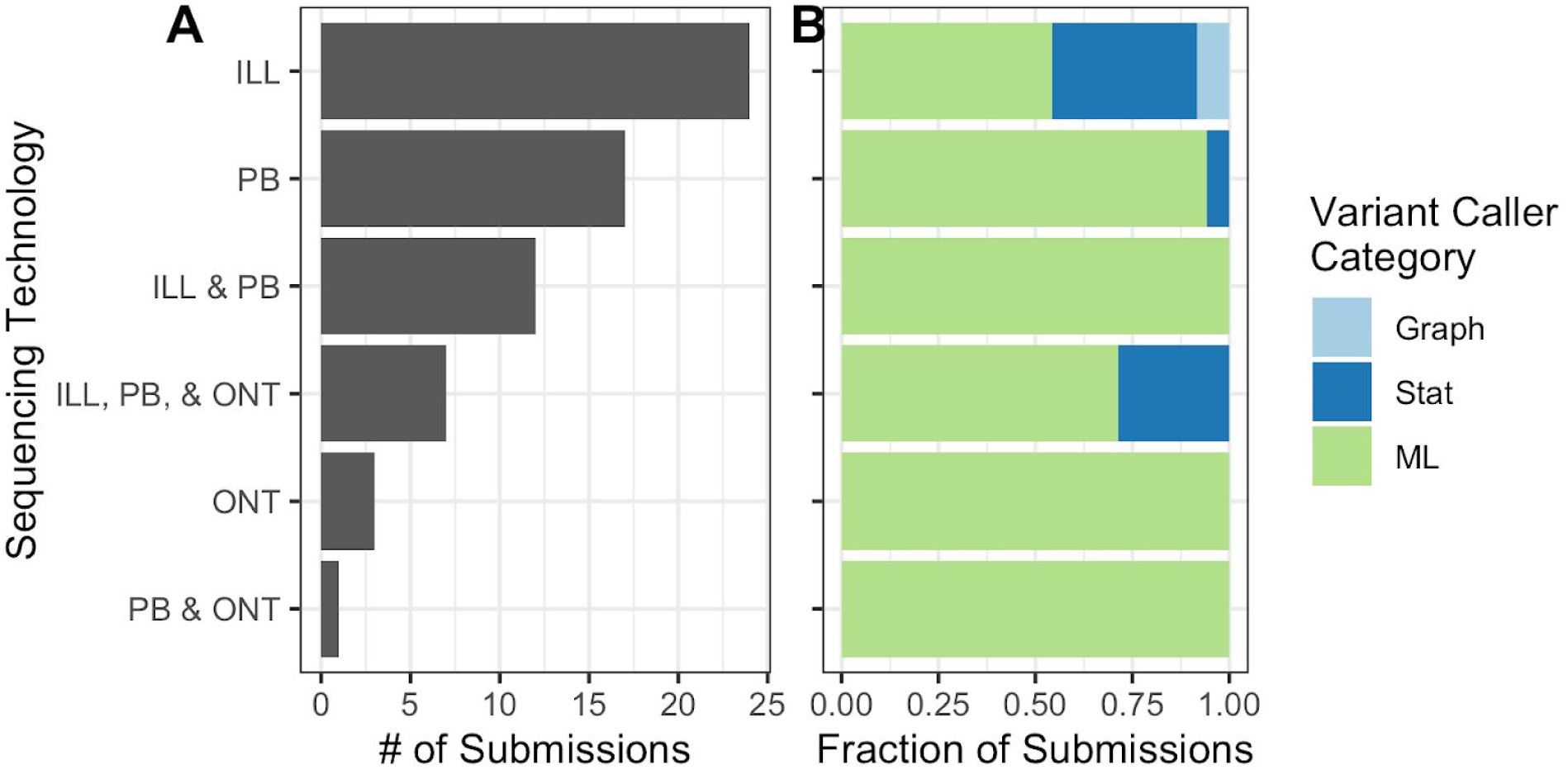
Challenge submission breakdown by (A) technology and (B) type of variant caller used.

Submissions were evaluated based on the averaged parents’ F1 scores for combined SNVs and INDELs. In all benchmark regions, the top performing submissions combined all technologies, followed by PacBio HiFi, Illumina, and ONT, with PacBio HiFi submissions having the best single-technology performance in each category (Fig. 3, Table 2). In contrast to all benchmark regions, submissions based on ONT performed better than Illumina in difficult-to-map regions despite ONT’s higher indel error rate. In fact, ONT-based variant calls had slightly higher F1 scores in difficult-to-map regions than in all benchmark regions, because the benchmark for difficult-to-map regions excludes homopolymers longer than 10 bp that are called by PCR-free short reads in easy-to-map regions. The best-performing short-read callsets (DRAGEN and Seven Bridges) used graph-based approaches, and the best-performing long-read callsets used ML (DeepVariant+PEPPER, NanoCaller, Sentieon, and Roche). Performance varied substantially across stratifications, with the best-performing multi-technology callsets having similar overall performance, although with error rates that varied by a factor of 10 in the Major Histocompatibility Complex (MHC).

**Table 2:** Summary of Challenge Top Performers. One winner was selected for each Technology/Genomic Region combination, and multiple winners were awarded in the case of ties. Winners were selected based on submission F1 score (SNV plus INDELs) for the blinded samples, HG003 and HG004.

| Technology | Genomic Region | Participant | F1 |
| --- | --- | --- | --- |
| MULTI | All Benchmark Regions* | Sentieon | 0.999 |
| MULTI | All Benchmark Regions* | Roche Sequencing Solutions | 0.999 |
| MULTI | All Benchmark Regions* | The Genomics Team in Google Health | 0.999 |
| MULTI | Difficult-to-Map Regions | Roche Sequencing Solutions | 0.994 |
| MULTI | MHC | Sentieon | 0.998 |
| ILLUMINA | All Benchmark Regions | DRAGEN | 0.997 |
| ILLUMINA | Difficult-to-Map Regions | DRAGEN | 0.969 |
| ILLUMINA | MHC | Seven Bridges Genomics | 0.992 |
| PACBIO | All Benchmark Regions | The Genomics Team in Google Health | 0.998 |
| PACBIO | Difficult-to-Map Regions | Sentieon | 0.993 |
| PACBIO | MHC | Sentieon | 0.995 |
| ONT | All Benchmark Regions | The UCSC CGL and Google Health | 0.965 |
| ONT | Difficult-to-Map Regions | The UCSC CGL and Google Health | 0.983 |
| ONT | MHC | Wang Genomics Lab | 0.972 |

\* Tied

**Figure 3:**
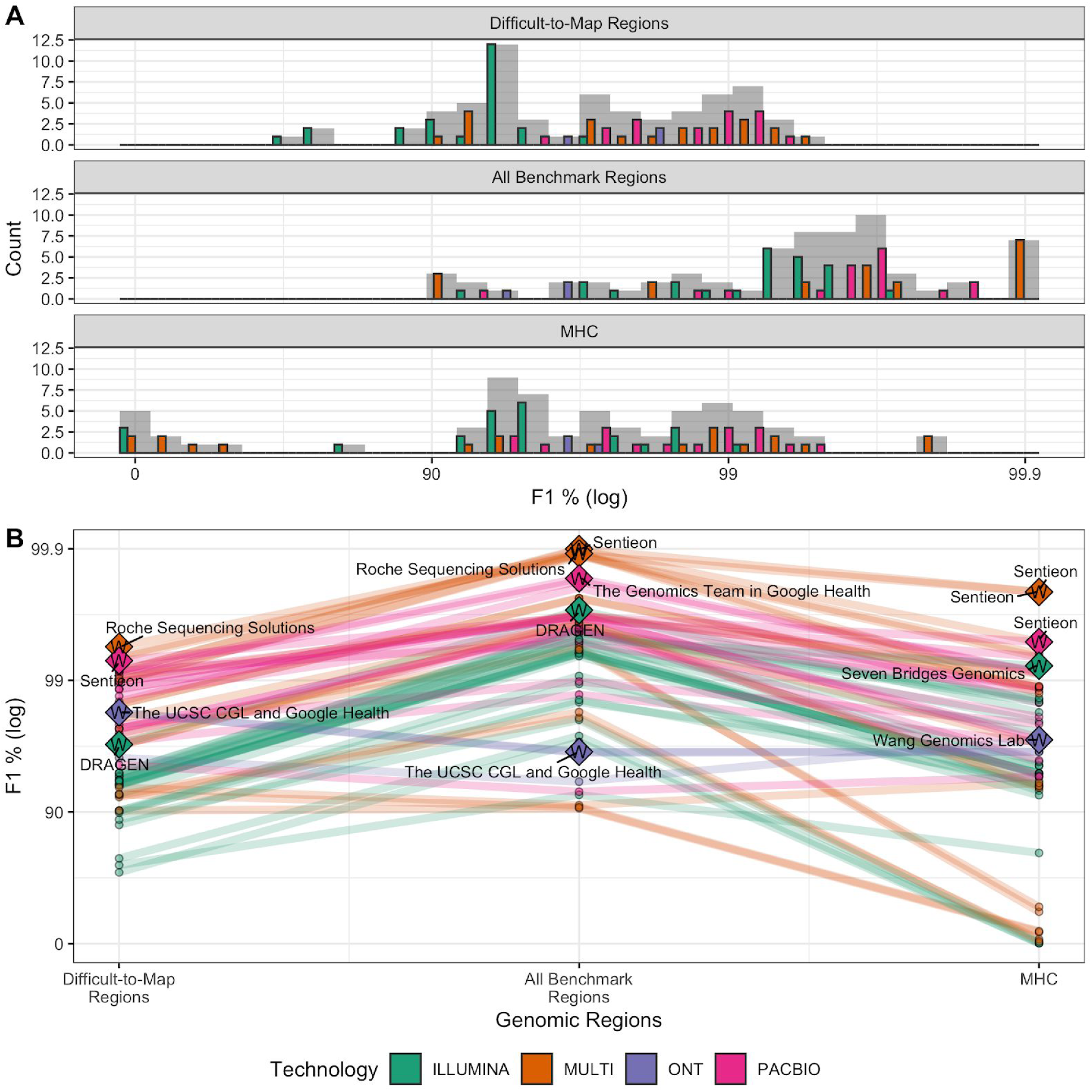
Overall performance (A) and submission rank (B) varied by technology and stratification (log scale). Generally, submissions that used multiple technologies (MULTI) outperformed single technology submissions for all three genomic context categories. Panel A shows a Histogram of F1 % (higher is better) for the three genomic stratifications evaluated. Submission counts across technologies are indicated by light grey bars and individual technologies by colored bars. Panel B shows individual submission performance. Data points represent submission performance for the three stratifications (difficult-to-map regions, all benchmark regions, MHC), and lines connect submissions. Category top performers are indicated by diamonds with “W”s and labeled with Team names.

### Challenge Highlights Innovations in Characterizing Clinically-important MHC

The medically relevant yet difficult to characterize MHC plays an important role in the immune response, for example recent research suggests HLA types encoded in the MHC plays a role in COVID severity (Nguyen et al., 2020). The MHC is a highly polymorphic ~5 Mb region of the genome that is particularly challenging for short-read methods (Fig. 4). In spite of difficulties associated with variant calling in this region, the Illumina graph-based pipeline developed by Seven Bridges (Rakocevic et al.) performed especially well in MHC (F1: 0.992). The Seven Bridges GRAF pipeline used in the Truth Challenge V2 utilizes a pan-genome graph that captures the genetic diversity of many populations around the world, resulting in a graph reference that accurately represents the highly polymorphic nature of the MHC region enabling improved read alignment and variant calling performance. The MHC region is more easily resolved with long-read based methods as these are more likely to map uniquely in the region. The ONT-NanoCaller Medaka (F1: 0.941) ensemble submission performed well on MHC, particularly for SNVs (F1: 0.992) and is the only method that performed as well in MHC as in all genomic benchmarking regions for SNVs. In general, submissions utilizing long-read sequencing data performed better than those only using short-read data. The difference in performance is less significant for INDELs than SNVs and likely due to differences in the sequencing error profile between the short- and long-read sequencing methods. INDELs were the dominant error type for both long-read sequencing methods and SNVs were the dominant error type for Illumina sequencing.

**Figure 4:**
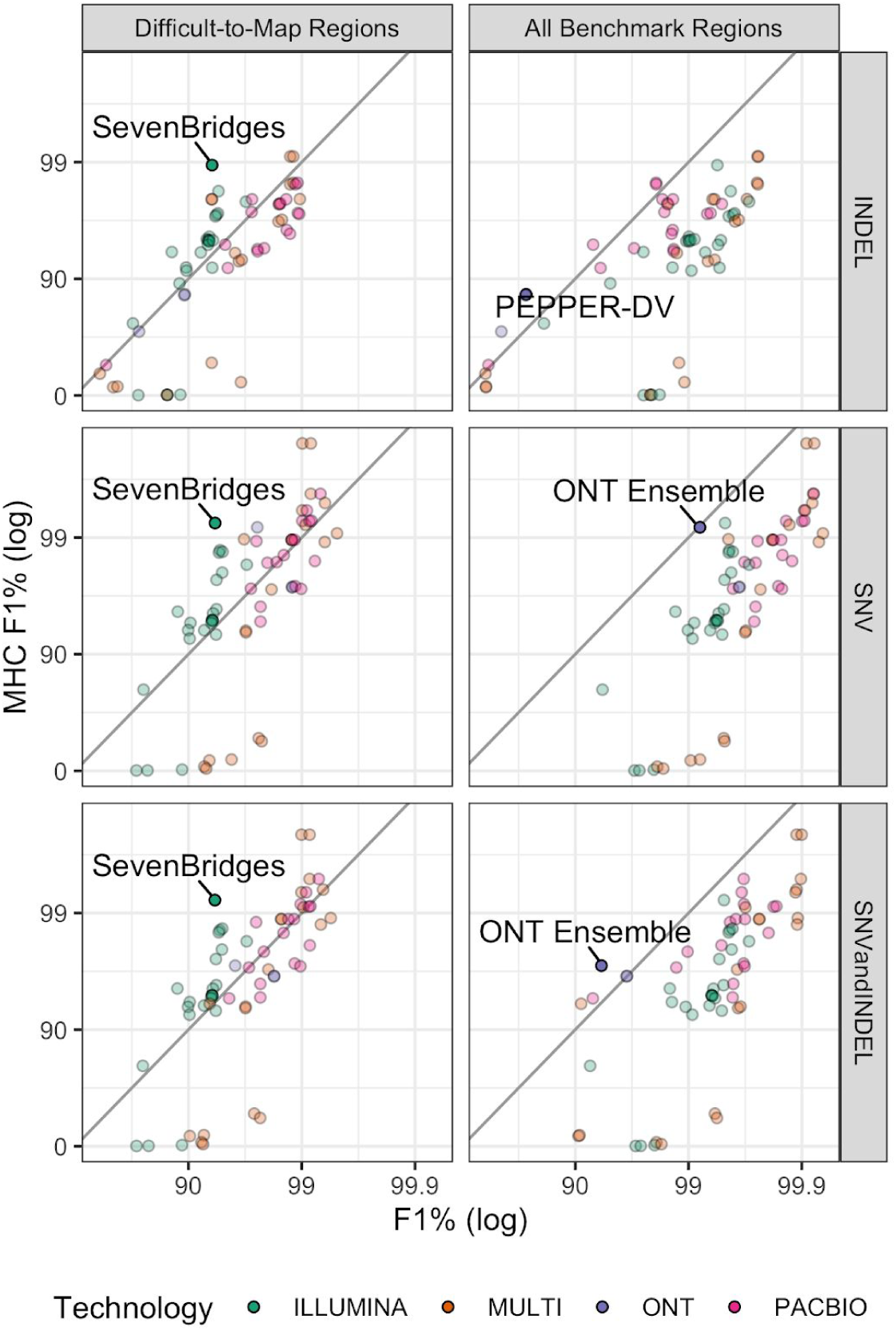
Submission performance comparison for F1 metric between MHC, all benchmark regions and difficult to map regions. Points above the diagonal black line perform better in MHC relative to all benchmark regions or the difficult to map regions. Submissions with the largest difference in performance between MHC and “Difficult-to-Map” or “All Benchmark Regions” for each subplot are labeled. SevenBridges - is a graph-based short read variant caller. ONT ensemble - is an ensemble of ONT variant callers NanoCaller, Clair and Medaka. PEPPER-DV - is the ONT PEPPER-DeepVariant haplotype-aware machine learning variant calling pipeline.

### Comparing performance for blinded and semi-blinded samples reveals possible over-tuning of some methods

Differences in performance were observed between the unblinded (HG002) and semi-blinded genomes (HG003 and HG004). We used the error rate ratio defined as the ratio of 1-F1 for the parents to the son (Eq. 1). The error rate ratio was generally larger for ML methods compared to non-ML methods and for long-read technologies compared to short-read technologies (Fig. 5). These error rate ratio differences are likely due to a combination of factors including differences in the sequence dataset characteristics between the three genomes, differences in the benchmark sets, and participants’ use of HG002 for model training and parameter optimization. Illumina variant call sets generally had smaller F1 score differences (median 1.06, range 0.98 - 4.38) regardless of the variant calling method used. The smaller error rate ratio may at least partially be due to the Illumina datasets being more consistent in coverage and base quality across the three genomes compared to the long-read datasets (Table 1, Fig. S1). For statistical-based variant callers such as GATK, differences in performance tend to be less than those among ML-based methods, especially for Illumina data. This smaller error rate ratio is likely due to the maturity of short-read variant calling compared to variant calling from long reads with ML-based variant callers. For the ONT-only variant callsets, the error rate ratio was less than 1, as the parents had higher F1 scores compared to the unblinded son. This is potentially due to the parents’ ONT datasets having higher coverage (85X) than the son’s (47X). The degree to which the ML models were over tuned to the training genome (HG002) and datasets as well as the impact of any over-tuning on variant calling accuracy warrants future investigation, but highlights the importance of transparently describing the training process. For the statistical-based variant callers, the observed drop in performance between the blinded and unblinded samples was likely due to optimizing algorithm parameters for HG002. Note that the parents do not represent fully blinded, orthogonal samples, since HG002 shares variants with at least one of the parents, and previous benchmarks were available for the easier regions of the parents’ genomes. These results highlight the need for multiple benchmark sets, sequencing datasets, and the value of established data types and variant calling pipelines.

**Figure 5:**
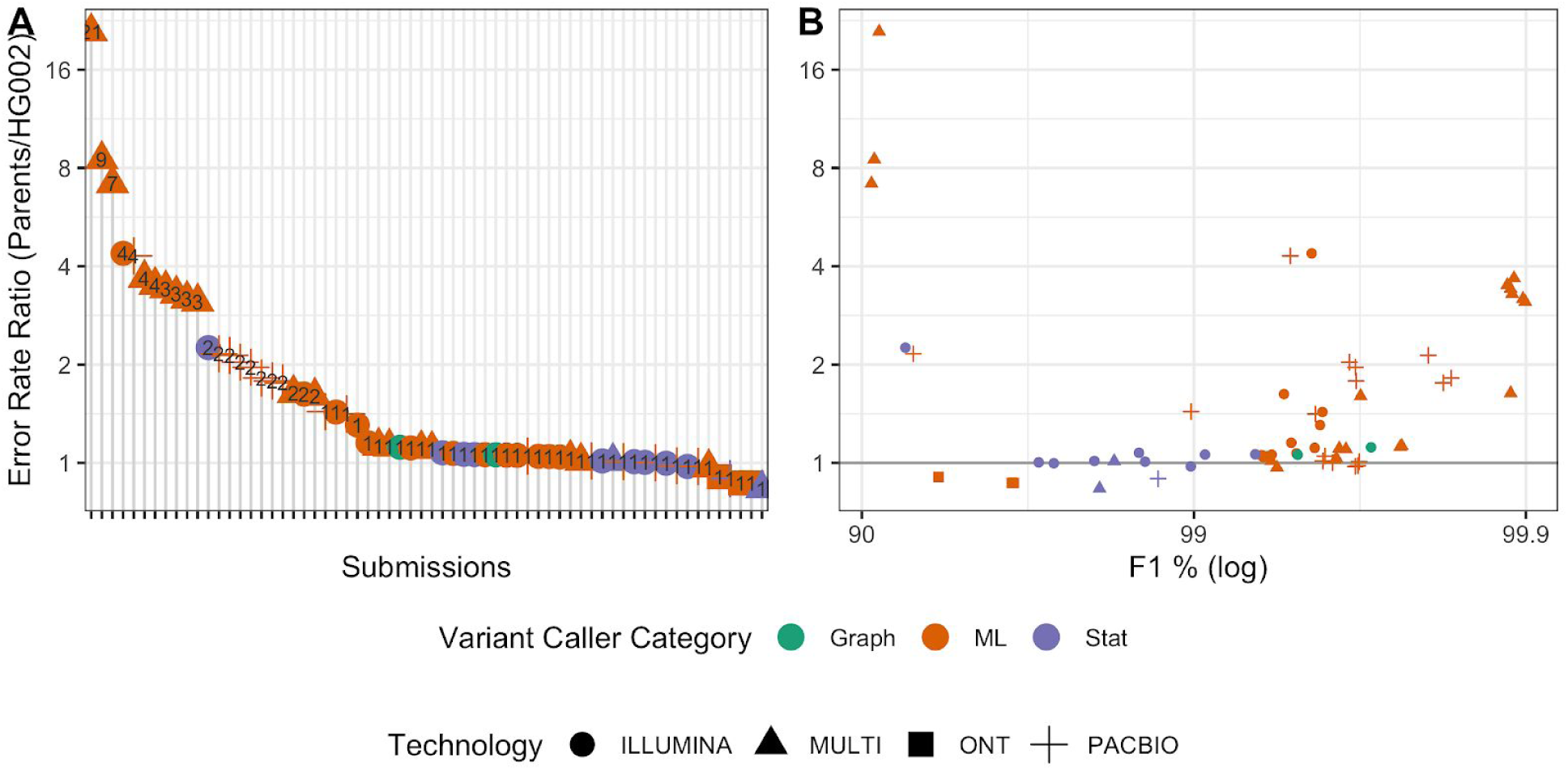
Ratio of error rates using semi-blinded parents’ benchmark vs. public son’s benchmark. (A) Submissions ranked by error rate ratio. (B) Comparison of error rate ratio to the overall performance for the parents (F1 in all benchmarking regions, as defined in eq. 1). Error rate defined as 1 - F1.

### Improved benchmark sets and stratifications reveal innovations in sequencing technologies and variant calling since the 2016 challenge

Since the first Truth Challenge held in 2016, variant calling, sequencing, and GIAB benchmark sets have substantially improved. The SNV error rates of the Truth Challenge V1 winners decrease by as much as 10-fold when benchmarked against the new V4.2 benchmark set, compared to the V3.2 benchmark set used to evaluate the first truth challenge (Fig. 6A). The V4.2 benchmark set covers 7% more of the genome than V3.2 (92% compared to 85% for HG002 on GRCh38), most importantly enabling robust performance assessment in difficult-to-map regions and the MHC (Wagner et al., 2020). The performance difference is more significant for SNVs compared to INDELs because the overall INDEL error rate is higher. Despite the higher coverage (50X) Illumina data used in the first challenge, several Illumina-only submissions from the V2 challenge performed better than all of the V1 challenge winners (Fig. 6B). This result highlights significant improvements in variant caller performance for short reads. Furthermore, advances in sequencing technologies have led to even higher accuracy, particularly in difficult-to-map regions. Improvements to the benchmarking set has allowed for more accurate variant benchmarking and, in turn, facilitated advances in variant calling methods, particularly ML-based methods which depend on the benchmark set for model training.

**Figure 6:**
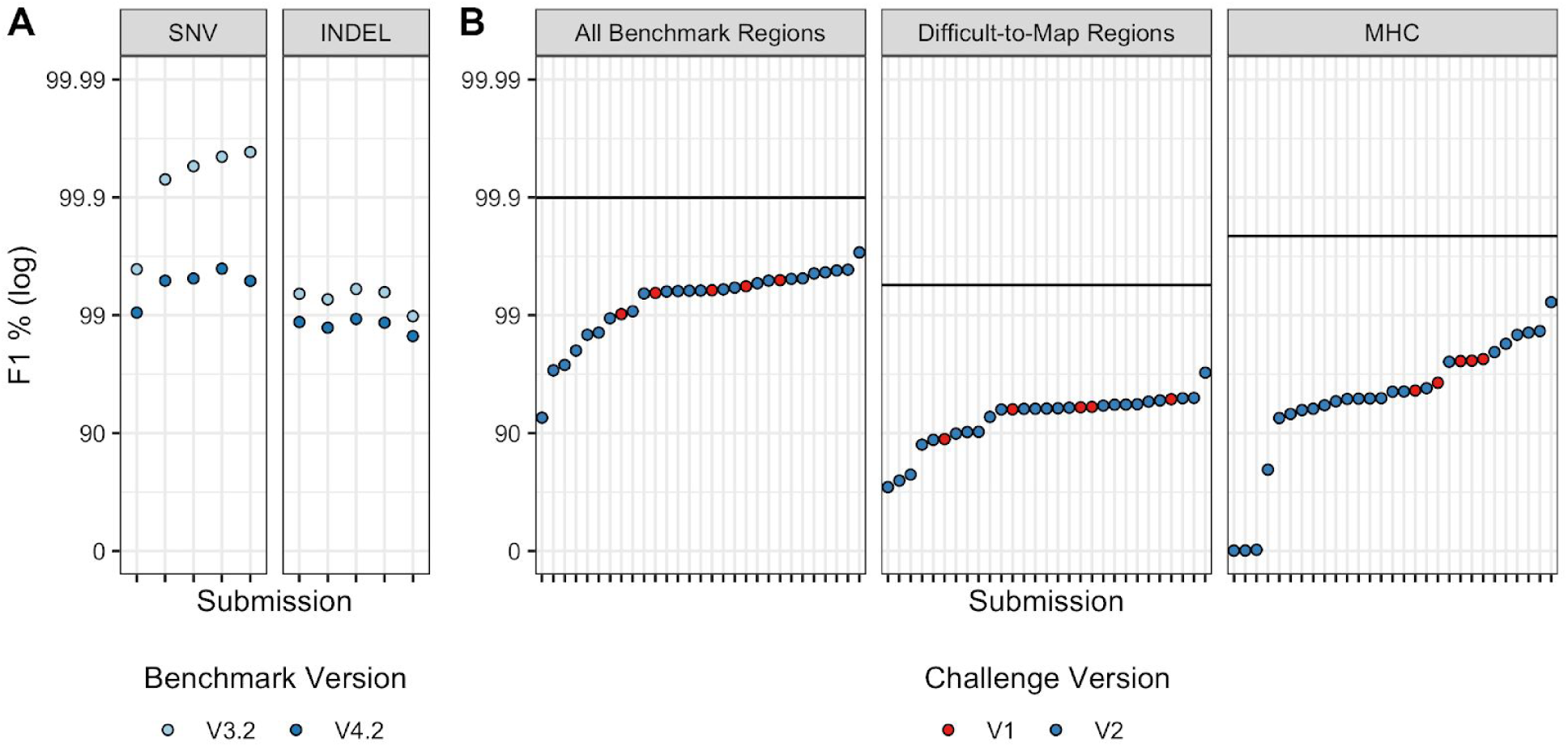
Comparison of benchmarking performance for (A) different benchmark sets and (B) challenges. (A) The 2016 (V1) Truth Challenge top performers F1 performance metric for SNVs and INDELs benchmarked against the V3.2 benchmark set (used to evaluate the first challenge) and V4.2 benchmark set (used to evaluate the second challenge). Performance metrics for the same variant calls decrease substantially vs. the V4.2 benchmark set because it includes more challenging regions. (B) Performance of V1 challenge top performers (using 50X Illumina sequencing) compared to V2 submissions (using only 35X Illumina sequencing) for the combined SNV and INDEL F1 metric and the V4.2 benchmark set used to evaluate the second truth challenge. The black horizontal lines represent the performance for the overall top performer, regardless of technology used, for each stratification. For the first challenge variant call sets for the blinded HG002 against GRCh37 were used to evaluate performance and for the second challenge variant calls for the semi-blinded HG003 and HG004 against GRCh38 were used to evaluate performance.

### New stratifications enable comparison of method strengths

As an example of the utility of stratifying performance in a more detailed way by genomic context with the new stratifications, we compared the ONT PEPPER-DeepVariant (ONT-PDV) submission to the Illumina DeepVariant (Ill-DV) submission (Fig. 7). The ONT-PDV submission has comparable overall performance to the Ill-DV submission for SNVs, providing an F1 of 99.64% and 99.57%, respectively, but performance differs >100-fold in some genomic context. Ill-DV SNV calls were more accurate in homopolymers and tandem repeats shorter than 200 bp in length. In contrast, ONT-PDV consistently had higher performance for segmental duplications, large tandem repeats, L1H, and other regions that are difficult to map with short reads. Due to the currently higher INDEL error rate for ONT R9.4 reads, Ill-DV INDEL variant calls are more accurate for nearly every genomic context, and the F1 for INDELs in all benchmark regions was 99.59% for Ill-DV compared to 72.54% for ONT-PDV. This type of analysis can help determine the appropriate method for a desired application, and understand how the strengths and weaknesses of technologies could be leveraged when combining technologies. High performing multi-technology submissions successfully incorporated callsets from multiple technologies by leveraging the additional coverage and complementary strengths of different technologies.

**Figure 7:**
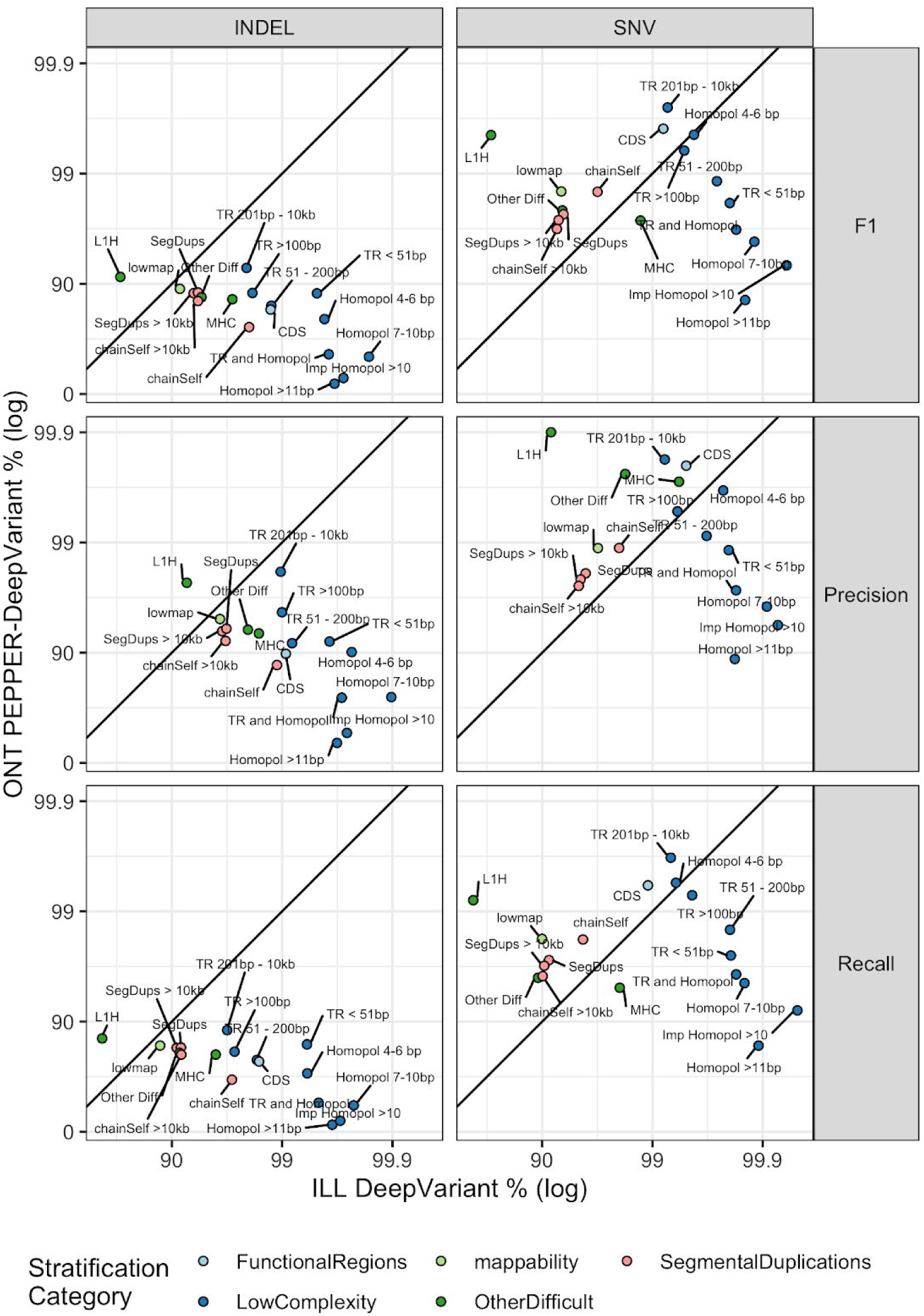
Comparison of ONT PEPPER-DeepVariant variant callset performance to Illumina DeepVariant by genomic context. Points above and below the diagonal line indicate stratifications where ONT PEPPER-DeepVariant submission performance metric was higher than the Illumina DeepVariant submission. The points are colored by stratification category.

## Discussion

Public genomics community challenges, such as the precisionFDA Truth Challenges described here, provide a public baseline for independent performance evaluation at a point in time against which future methods can be compared. It is important to recognize the advancements and limitations of the benchmarks used in these challenges. For example, the GIAB V3.2 benchmark set used to evaluate the first precisionFDA Truth Challenge submissions only included the easier regions of the genome (https://precision.fda.gov/challenges/truth/results), excluding most segmental duplications and difficult-to-map regions, as well as the highly polymorphic MHC. This is evidenced by the fact that when the first Truth Challenge winners were benchmarked against the new V4.2 benchmark set, which included more difficult regions of the genome, the performance metrics decreased as much as 10-fold (Fig. 6A). It is important to note that these challenges are not just to compare and inspire new methods, but to give the research and clinical sequencing community insight into what is currently possible in terms of accuracy and which methods might be applicable to the experiment in mind.

Public community challenges further help drive the methods development. A number of ground-breaking mapping+variant calling pipelines were developed, optimized, and made available as part of this challenge. For example, the new experimental DRAGEN method used graph-based mapping and improved statistical variant calling approaches to call variants in segmental duplications and other regions previously difficult to map with short reads. DRAGEN’s graph-based mapping method used alt-aware mapping for population haplotypes stitched into the reference with known alignments, effectively establishing alternate graph paths that reads could seed-map and align to. This reduced mapping ambiguity because reads containing population variants were attracted to the specific regions where those variants were observed. The Seven Bridges GRAF pipeline used genome graphs to align reads and call variants in the whole genome. The graph reference is constructed by augmenting the linear genome reference with existing genetic information. A pan-genome graph capturing the genetic diversity of many populations around the world was used by the Seven Bridges team for the challenge. The pan-genome graph was constructed by incorporating multiple variant databases (such as 1000 Genomes, Simons Diversity Project, gnomAD) and also relocating the alt-contigs in the GRCh38 assembly to their canonical positions as alternate haplotypes/edges on the graph reference. This results in a graph reference that can accurately represent, for instance, the highly polymorphic nature of the MHC region and therefore enable improved read alignment and variant calling performance in this region by the Seven Bridges GRAF pipeline.

For the long read methods, innovative machine learning-based methods were developed for this challenge. The PEPPER-DeepVariant used new approaches for selecting candidate variants and called genotypes accurately for small variants despite the relatively high error rate in raw ONT reads. Several new ML methods enabled highly accurate variant calling from the new PacBio HiFi technology. While different sequencing technologies have different strengths, robust integration of data from different technologies is challenging. Several submissions used new approaches to integrate multiple technologies and leverage the independent technology-specific information as well as additional coverage from the combining data to perform better than any individual technology.

Along with the new benchmark set and sequencing data types, we used new genomic stratifications to evaluate submission performance in different contexts, highlighting methods that performed best in particularly challenging regions. For example, the Seven Bridges GRAF Illumina and NanoCaller ONT submissions performed particularly well in the MHC, and the Sentieon PacBio HiFi submission performed particularly well in both the MHC and difficult-to-map regions. These submissions might have been overlooked if the performance was not stratified by context. The new stratifications presented here represent a valuable resource to the community for use in evaluating and optimizing variant calling methods. Stratifying performance by genomic context can be valuable in at least three ways including 1) assessing the strengths and weaknesses of a method for different genome contexts and variant types, which is, for example, critical in clinical validation of bioinformatics methods (Roy et al., 2018); 2) aide in understanding which variants are not assessed by the benchmark; and 3) aide in selecting the technology and bioinformatics methods that are best suited for the genomic regions of interest, e.g. MHC.

Deep learning and ML have advanced variant calling, particularly by enabling faster adoption of new sequencing technologies. In this context, care should be taken to evaluate over-training and be transparent about the data used for training, tuning and testing. Based on results from this challenge, there is likely at least some overfitting to training samples. Overtraining can occur both to the individual (HG002) and to the properties of the particular sequencing runs that are used for training. Non-ML methods can also overfit, because coding and parameter selection will be guided by performance on the development set. For example, short-read variant callers that use information from long-read sequencing datasets may perform better for samples or populations included in the long-read data. Similarly, methods using graph references may perform better for samples or populations used in constructing the graph. Having clear provenance of training samples including multiple ethnicities and regions is important for the field. These results also highlight the importance of developing additional genomically diverse benchmark sets.

This challenge spurred the development and public dissemination of a diverse set of new bioinformatics methods for multiple technologies. It provides a public resource for capturing method performance at a point in time, against which future methods can be compared. New versions of these methods and new methods will continue to improve upon the methods presented here. For example, immediately after the challenge, two different participants combined the strengths of a new mapping method for long reads from one submission (winnowmap) with a new variant calling method from another submission (PEPPER-DeepVariant) to get improved results (Fig. S3)(Jain et al., 2020). The GIAB benchmarks help enable the ongoing improvements, and GIAB/GA4GH benchmarking tools enable identification of strengths and weaknesses of any method in stratified genome contexts. The new variant calling methods presented in this challenge can help improve future versions of benchmarks that will be critical as variant calling methods and sequencing technologies continue to improve, thus driving the advancement of research and clinical sequencing.

## Supporting information

Submission Methods

## Acknowledgements

The authors would like to thank the anonymous challenge participants, as well as Drs. Megan Cleveland and Hua-Jun He for feedback on the manuscript. Certain commercial equipment, instruments, or materials are identified in this paper in order to specify the experimental procedure adequately. Such identification is not intended to imply recommendation or endorsement by NIST, nor is it intended to imply that the materials or equipment identified are necessarily the best available for the purpose.

## Funding

Ministerio de Ciencia e Innovación (RTC-2017-6471-1; AEI/FEDER, UE) co-financed by the European Regional Development Funds ‘A way of making Europe’ from the European Union and Cabildo Insular de Tenerife (CGIEU0000219140) to C.F.. FJS was supported by NIH (UM1 HG008898) UA, QL and KW were supported by NIH (GM132713).

## Author Contributions

J.W., J.M.Z., N.D.O., E.J., E.B., S.H.S., A.P., E.M., O.S., S.W., and F.J.S. contributed to challenge design. E.J., E.B., S.H.S., A.P., E.M., O.S., and S.W. coordinated the challenge. D.J., J.M.L., A.M., L.A.R., C.F., K.K, A.M., K.S., T.P., M.J., B.P., P.C., A.K., M.N., G.B., S.G., H.Y., A.C., R.E., M.B., G.B., G.L., C.M., L.T., Y.D., S.Z., J.M., R.T., G.P., J.T., C.B., S.D., D.K., D.T., Ö.K., G.B., K.N., E.A., R.B., I.J.J., A.D., V.S., A.J., H.S.T., V.J., M.R., B.L., C.R., S.C., R.M., M.U.A., Q.L., K.W., S.M.E.S., L.T., M.M., C.H., C.J., H.F., Z.L., and L.C. participated in the challenge and contributed to the methods section. N.D.O., J.M., J.W. and J.W.Z. contributed to the data curation and analysis. All authors reviewed and edited the manuscript.

## Declaration of Interests

C.B. is an employee and shareholder of SAGA Diagnostics AB.

A.C., P.C., A.K., M.N., G.B., S.G., and H.Y. are employees of Google and A.C. is a shareholder

S.D., D.K., D.T., Ö.K., G.B., K.N., E.A., R.B., I.J.J., A.D., V.S., A.J., and H.S.T. are employees of Seven Bridges Genomics

O.S. and S.T.W. are employees of DNAnexus

G.L, C.M, L.T., Y.D., and S.Z. are employees of Genetalks

V.J., M.R., B.L., C.R, S.C., and R.M are employees of Illumina

S.M.E.S., and M.M. are employees of Roche

C.H. is an employee of Wasai Technology

H.F., Z.L, and L.C. are employees of Sentieon Inc.

## Materials availability

DNA extracted from a single large batch of cells for each genome is publicly available in National Institute of Standards and Technology Reference Material 8392. Cell lines from which these DNA are extracted are publicly available as GM24385 (RRID:CVCL_1C78), GM24149 (RRID:CVCL_1C54), and GM24143 (RRID:CVCL_1C48) at the Coriell Institute for Medical Research National Institute for General Medical Sciences cell line repository.

## Data and Code Availability

Input sequencing data, participant submitted VCFs, and benchmarking results are available at https://doi.org/10.18434/mds2-2336. Sequencing data are available on the precisionFDA platform and SRA, see Supplemental Material and methods for additional information. Code used to analyze challenge results presented in the manuscript and benchmarking results files are available at https://github.com/usnistgov/giab-pFDA-2nd-challenge.

## Methods

The samples were sequenced under similar sequencing conditions and instruments across the three genomes. For the Illumina dataset, 2×151 bp high coverage PCR-free library was sequenced on the NovaSeq 6000 System (manuscript in-prep). The datasets were downsampled to 35X based on recommended coverage used in variant calling. The full 50X datasets were downsampled to 35X using seqtk (https://github.com/lh3/seqtk) and the following command seqtk sample - s100 {fastq} 0.752733763. For PacBio HiFi, we used the library size and coverage recommended at the time by PacBio for variant calling, ~35X 15 kb libraries. For HG002, 4 SMRT Cells were sequenced using the Sequel II System with 2.0 chemistry. Consensus basecalling was performed using the “Circular Consensus Sequencing” analysis in SMRT Link v8.0, ccs version 4.0.0. Data from the 15 kb library SMRT Cells were merged and downsampled to 35X. The combined flowcell FASTQs were downsampled using seqtk (v1.3r106, https://github.com/lh3/seqtk) to a median coverage across chromosomes 1 to 22 of 35X. Coverage was verified by mapping reads to GRCh38 using minimap2 (Li, 2018) and coverage was calculated with mosdepth v0.2.9 (Pedersen and Quinlan, 2018) using a window size of 10 kb. The PacBio HiFi data are available on SRA under the following BioProjects; HG002 - PRJNA586863, HG003 - PRJNA626365, and HG004 - PRJNA626366. The ONT dataset was generated using the unsheared DNA library prep, methods described elsewhere (Shafin et al., 2020), and consisted of pooled sequencing data from three PromethION R9.4 flowcells. Basecalling was performed using Guppy Version 3.6 (https://community.nanoporetech.com). Data from three ONT PromethION flow cells were used for each of the 3 genomes, but the resulting coverage was substantially higher for the parents (85X) than the child (47X) with similar read length distributions (Fig. S2). See Supplemental Material for links to the FASTQ files provided to participants on the precisionFDA platform.

The HG002 V4.1 benchmark set was unblinded and available to participants for model training and methods development. We used the blinded HG003 and HG004 V4.2 benchmark sets to evaluate performance. The V4.1 and V4.2 benchmark sets are the latest versions of the GIAB small variant benchmark set, which utilize long- and linked-read sequencing data to expand the benchmark set into difficult regions of the genome (Wagner et al., 2020). Prior to submission, participants could benchmark their HG002 variant callsets using the precisionFDA comparator tool (https://precision.fda.gov/apps/app-F5YXbp80PBYFP059656gYxXQ-1, a free precisionFDA account is required for access). The comparator tool is an implementation of the GA4GH small variant benchmarking tool hap.py (Krusche et al., 2019, https://github.com/Illumina/hap.py) with vcfeval (Cleary et al., 2015) on the precisionFDA platform. The same comparator tool was used to evaluate submission performance against the HG003 and HG004 V4.2 benchmark sets. To evaluate performance for different genomic contexts, the V2.0 genome stratifications were used (https://data.nist.gov/od/id/mds2-2190). Submissions were evaluated using the geometric mean of the HG003 and HG004 combined SNVs and INDELs F1 scores (eq. 1). We use the error rate ratio (ERR), defined as the ratio of 1-F1 for the parents to the son.

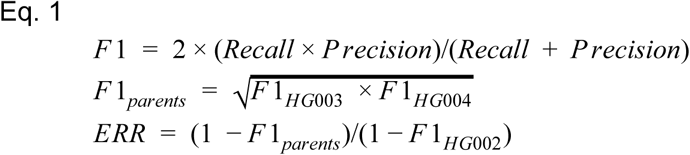

The V2.0 genome stratifications are an update to the GA4GH genomic stratifications utilized by hap.py (Krusche et al., 2019). The V2.0 stratifications are a pared down set of stratifications with improved strata for complex regions, such as tandem repeats and segmental duplications, as well as new genome-specific stratifications for suspected copy number variants (CNVs) and known errors in the reference genome (Table 3). The GRCh38 V2.0 stratifications includes 127 stratifications. See supplemental material for a detailed description of the genomic stratifications.

**Table 3:** Summary table of the V2.0 GIAB genome stratifications. The new stratification set includes the union of multiple stratifications as well as “not in” stratifications, which are useful in evaluating performance outside specific difficult genomic contexts.

| Stratification Group | Description | # Strats | Example Stratifications | Useful for |
| --- | --- | --- | --- | --- |
| FunctionalRegions | Coding regions | 2 | CDS, not in CDS | Evaluating performance in coding regions more likely to be functional |
| GC-content | Various ranges of GC-content | 14 | GC < 25%; 30% < GC < 55% | identifying GC bias in variant calling performance |
| Low Complexity |  | 22 |  | evaluating performance in locally repetitive, difficult to sequence contexts |
| - Homopolymers | Identification of homopolymers by length | 4 | Homopolymers > 101 bp; imperfect homopolymers > 10 bp | evaluating performance in homopolymers, where systematic sequencing errors and complex variants frequently occur |
| - Simple Repeats | Di, tri, and quad-nucleotide repeats of different lengths | 9 | Di-nucleotide repeats 11-50 bp; di-nucleotide repeats > 200 bp | evaluating performance in exact Short Tandem Repeats where systematic sequencing errors and complex variants frequently occur, and variant calls are challenging if the read length is insufficient to traverse the entire repeat |
| - Tandem Repeats | Tandem repeats of different lengths | 5 | Tandem repeats between 51 - 200 bp; tandem repeats > 10 kb | evaluating performance in exact Short Tandem Repeats and Variable Number Tandem Repeats where systematic sequencing errors and complex variants frequently occur, and variant calls are challenging if the read length is insufficient to traverse the entire repeat |
| Other Difficult | Various difficult regions of the genome | 6 | MHC; VDJ | evaluating performance in or excluding regions where variants are difficult to call and represent due to limitations of the reference genome (e.g. gaps or errors) or being highly polymorphic in the population (MHC). |
| Segmental Duplications | Segmental duplications defined using multiple methods and limited to segdups > 10kb | 9 | Segdups > 10 kb; selfChain | Regions with multiple similar copies in the reference, making them challenging to map and assemble. |
| Genome Specific | Difficult regions of the genome specific to one or more of the GIAB genomes. Including but not limited to complex variants, copy number variants, and structural variants. | 65 | CNVs, complex variants | evaluating performance in or excluding regions in each GIAB reference sample where small variants can be challenging to call (e.g., complex variants) or represent (e.g., CNVs and SVs) |

Participant-provided variant calling methods are included as Supplemental Table 1. Fifteen of the twenty participants, including all the challenge winners, provided methods to be made publicly available for this manuscript, a requirement for co-authorship. A random unique identifier was generated for every submission. For participants intending to remain anonymous, the unique identifier was used as the participant and submission names in the methods description.

To better understand how improvements in variant calling methods, sequencing technologies, and benchmark sets affect performance metrics, we benchmarked the first challenge winners against the new benchmark. For the first challenge, participants submitted variant calls for HG001 and HG002 against GRCh37 using Illumina short-read sequencing data, 2×150 bp 50X coverage (higher than the more commonly used 35X in the V2 Challenge). We benchmarked the winners of the first challenge (https://precision.fda.gov/challenges/truth/results) against the V4.2 HG002 GRCh37 benchmark set. The performance metrics for the V3.2 benchmark set were obtained from the precisionFDA challenge website.

The input benchmarking results and code used to perform the analyses presented below are available (https://github.com/usnistgov/giab-pFDA-2nd-challenge). The statistical programming language R was used for data analysis. Rmarkdown was used to generate individual results (Xie et al., 2020). Packages in the Tidyverse were used for data manipulation and plotting specifically ggplot, tidyr, and dplyr (Wickham et al., 2019).

## Supplemental Figures

**Figure S1:**
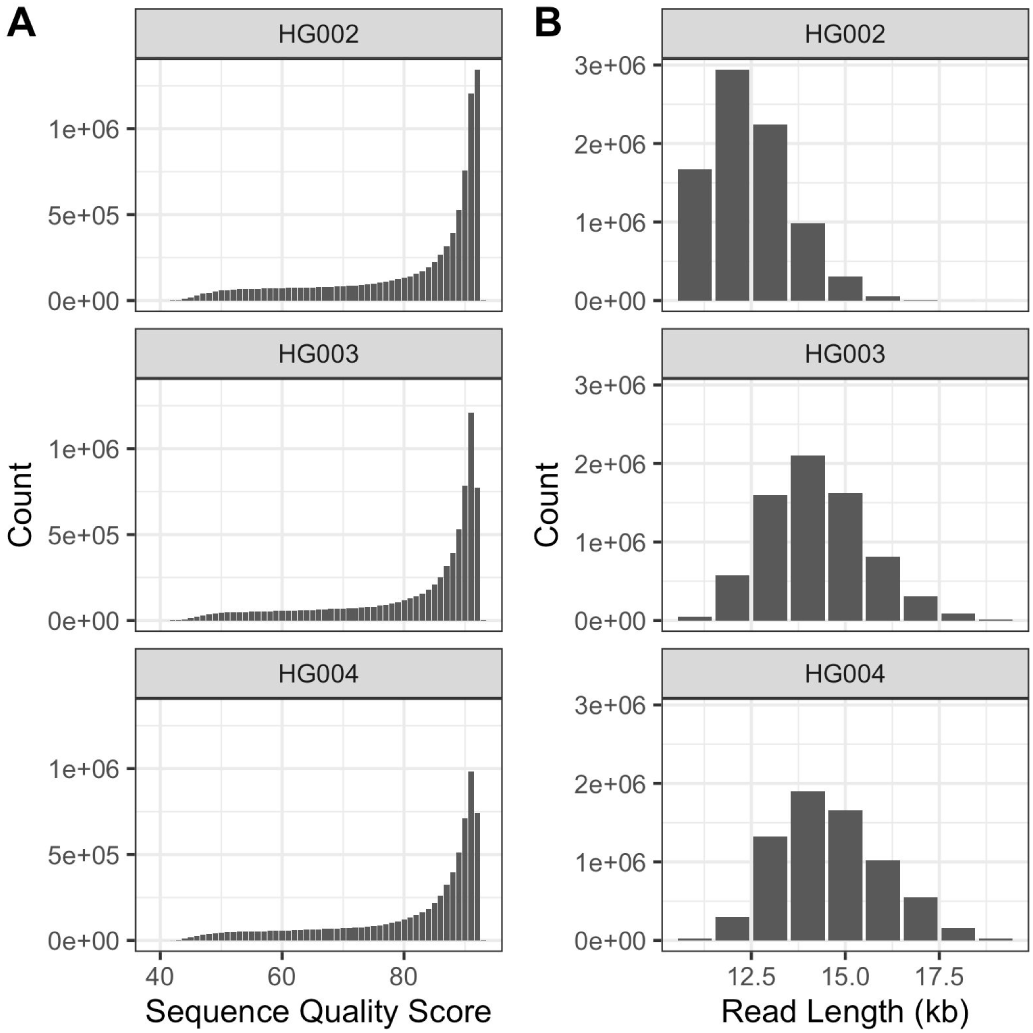
Read length and sequence quality score distributions for the three PacBio HiFi datasets. Sequence data metrics were calculated using FastQC.

**Figure S2:**
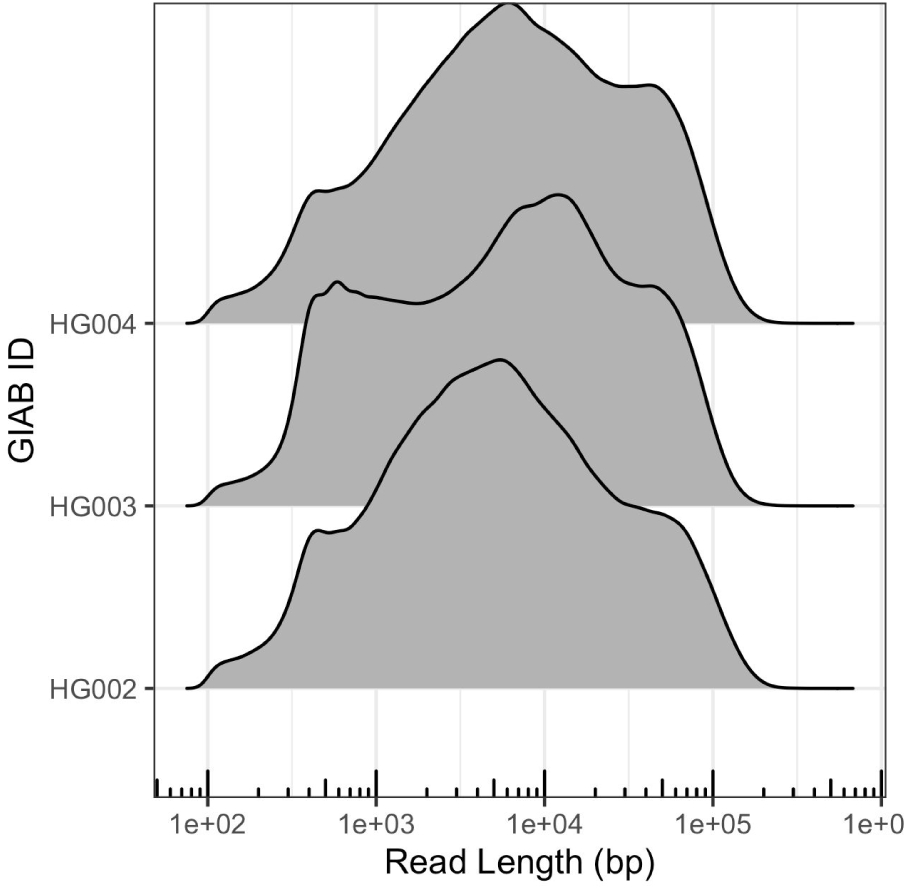
Read length distribution for the three ONT PromethION datasets.

**Figure S3:**
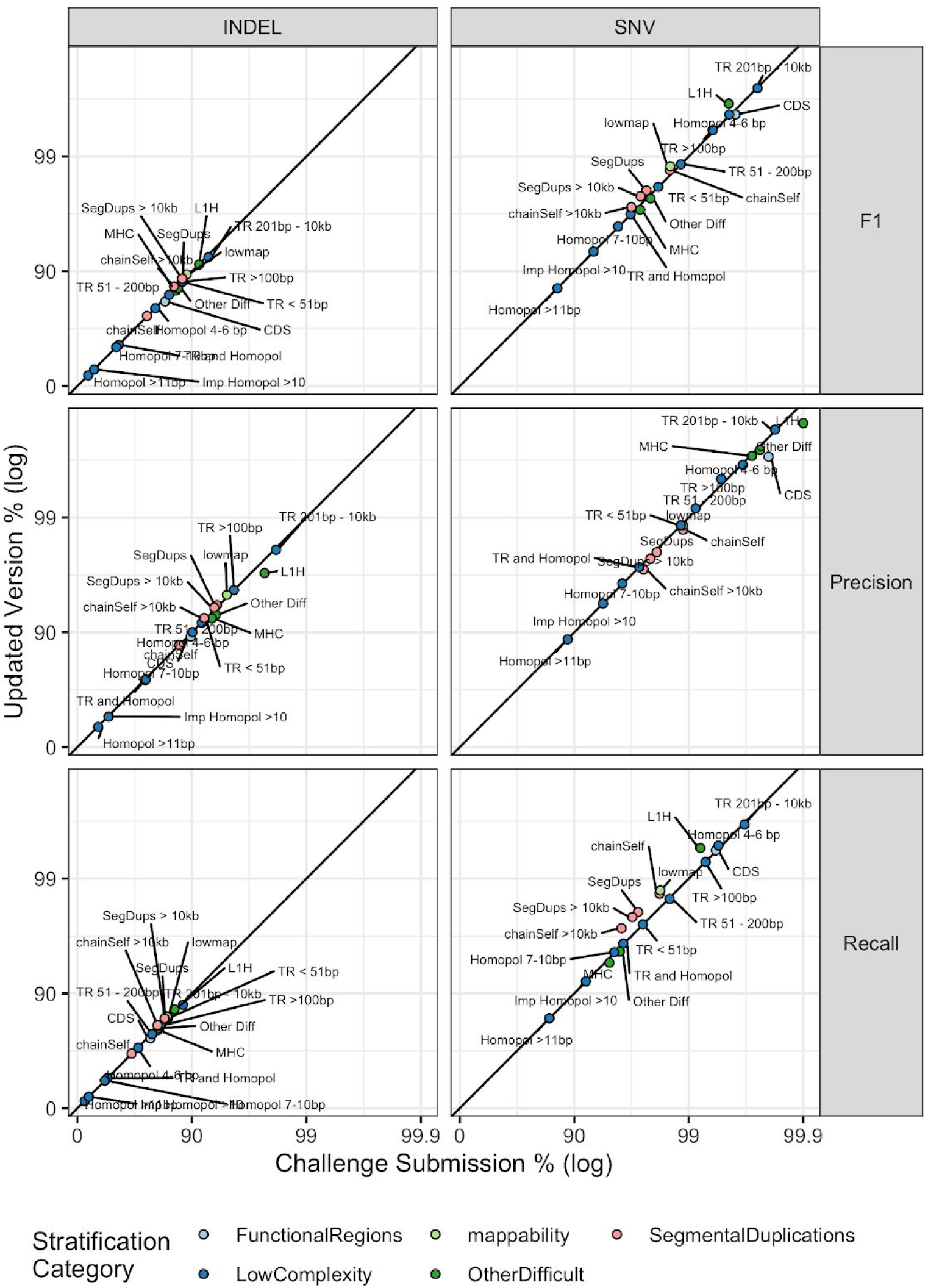
Comparison of submitted version of the ONT PEPPER-DeepVariant variant callset performance to an updated version. After the challenge ended a new mapping algorithm for long read data, winnowmap, was released. Winnowmap uses weighted minimizers to improve read mapping in repetitive genomic regions (https://doi.org/10.1093/bioinformatics/btaa435). The updated variant callset utilizes this new read mapping algorithm in its pipeline. Points above and below the diagonal line indicate stratifications where the updated callset performance metric was higher than the challenge submission. The points are colored by stratification category.

## Supplemental Material

### Challenge Sequencing Datasets

Links to FASTQ files provided to challenge participants on the precisionFDA platform. A free precisionFDA account is required for file access.

HG002 (NA24385)

- Illumina
  - precisionFDA:
    - [HG002.novaseq.pcr-free.35x.R1.fastq.gz](https://precision.fda.gov/files/file-FpZG9Jj0xbJy0QXjB7yb6fpX-1)
    - [HG002.novaseq.pcr-free.35x.R2.fastq.gz](https://precision.fda.gov/files/file-FpZG9Jj0xbJfGbf01qBgKyg1-1)
- PacBio HiFi
  - precisionFDA: [HG002_35x_PacBio_14kb-15kb.fastq.gz](https://precision.fda.gov/files/file-Fpx3BKj0pzk1xGyb9FjZYZ0x-1)
  - SRA:
    - Bioproject: PRJNA586863
    - Accessions: SRX7083054, SRX7083055, SRX7083056, and SRX7083057
- Oxford Nanopore
  - precisionFDA: [HG002_GM24385_1_2_3_Guppy_3.6.0_prom.fastq.gz](https://precision.fda.gov/files/file-Fpk3KGj0fQX2VjXjF8YXqz93-1)

HG003 (NA24149)

- Illumina
  - precisionFDA:
    - [HG003.novaseq.pcr-free.35x.R1.fastq.gz](https://precision.fda.gov/files/file-FpZBpZQ0xbJbqX5fGjgz9B1G-1)
    - [HG003.novaseq.pcr-free.35x.R2.fastq.gz](https://precision.fda.gov/files/file-FpZBpZQ0xbJYF2yfG8Y9kJ15-1)
- PacBio HiFi
  - precisionFDA: [HG003_35x_PacBio_14kb-15kb.fastq.gz](https://precision.fda.gov/files/file-FpvKx6j0Fx382vPq0p54Vk23-1)
  - SRA
    - Bioproject Accession: PRJNA626365
    - SRA Accessions: SRX8136474, SRX8136475, SRX8136476, and SRX8136477
- Oxford Nanopore
  - precisionFDA: [HG003_GM24149_1_2_3_Guppy_3.6.0_prom.fastq.gz](https://precision.fda.gov/files/file-Fpk3VVQ0fQX12Pvb925XP843-1)

HG004 (NA24143)

- Illumina
  - precisionFDA:
    - [HG004.novaseq.pcr-free.35x.R1.fastq.gz](https://precision.fda.gov/files/file-FpZG9Jj0xbJQ40QFGjPf019X-1)
    - [HG004.novaseq.pcr-free.35x.R2.fastq.gz](https://precision.fda.gov/files/file-FpZG9Jj0xbJpY09b1vfvgbbv-1)
- PacBio HiFi
  - precisionFDA: [HG004_35x_PacBio_14kb-15kb.fastq.gz](https://precision.fda.gov/files/file-FpvKxv80vzGJ1xQK9GX5qG4k-1)
  - SRA
    - Bioproject Accession: PRJNA626366
    - SRA Accessions: SRX8137018, SRX8137019, SRX8137020, and SRX8137021
- Oxford Nanopore
  - precisionFDA: [HG004_GM24143_1_2_3_Guppy_3.6.0_prom.fastq.gz](https://precision.fda.gov/files/file-Fpk3VZQ0fQXKP8xk381VGYvg-1)

## Genome Stratifications

The Global Alliance for Genomics and Health (GA4GH) Benchmarking Team and the Genome in a Bottle (GIAB) Consortium v2.0 stratification BED files are intended as standard resource of BED files for use in stratifying true positive, false positive, and false negative variant calls. The stratification BED files can be accessed from the NIST Public Data Repository, https://data.nist.gov/od/id/mds2-2190. All stratifications that utilize the GRCh38 reference use the reference without decoy or ALT loci (ftp://ftp.ncbi.nlm.nih.gov/genomes/all/GCA/000/001/405/GCA_000001405.15_GRCh38/seqs_for_alignment_pipelines.ucsc_ids/GCA_000001405.15_GRCh38_no_alt_analysis_set.fna.gz, link checked 08/31/2020).

### Functional Regions

Two Functional Region stratifications were created to stratify variants inside and outside of coding regions. The coding regions were extracted from the RefSeq GFF file (https://ftp.ncbi.nlm.nih.gov/genomes/all/GCF/000/001/405/GCF_000001405.39_GRCh38.p13/GCF_000001405.39_GRCh38.p13_genomic.gff.gz, link checked 08/31/2020). Non-overlapping complement regions for some stratifications are also provided, as “notin” files.

### GC Content

Fourteen GC content stratifications were created to stratify variants into different ranges of GC content. Using the seqtk algorithm (https://github.com/lh3/seqtk, link checked 08/31/20) with the GRCh38 reference, >=x bp regions with >y% or <y% GC were identified. The output was further processed to generate 100 bp ranges of GC with an additional 50 bp slop on either side (Ross et al., 2013).

Note that after adding 50 bp slop, 274,889 bp overlap between gc30 and gc65, or 0.05% of gc30 and 0.5% of gc65, or 0.07% of gc30 and 0.5% of gc65. The BED files with different GC ranges are almost exclusive of each other, but not completely.

We chose to stratify regions with <30% or >55% GC because these regions had decreased coverage or higher error rates for at least one of the technologies in Ross et al. (2013), and we added 55-60 and 60-65 because we found increased error rates in these tranches in exploratory work.

### Genome Specific

For each GIAB genome, Genome Specific stratifications were created to identify variants in difficult regions due to potentially difficult variation in the NIST/GIAB sample, including (1) regions containing putative compound heterozygous variants, (2) regions containing multiple variants within 50 bp of each other, (3) regions with potential structural variation and copy number variation. GRCh37 stratifications were generated using vcflib vcfgeno2haplo and Unix commands to identify complex and compound variants in benchmark VCF files from GIAB (Zook et al., 2019) for all samples, as well as Platinum Genomes (Eberle et al., 2017), and Real Time Genomics (Cleary et al., 2014) for HG001/NA12878. To generate GRCh38 Genome Specific stratifications, the GRCh37 Genome Specific complex/compound/SVs BED files were remapped to GRCh38 using the NCBI Remapping Service (https://www.ncbi.nlm.nih.gov/genome/tools/remap). Non-overlapping complement regions for some stratifications are also provided, as “notin” files.

### Functional Technically Difficult

The Functional Technically Difficult stratification is used in stratifying variants by different functional, or potentially functional, regions that are also likely to be technically difficult to sequence. A list of GRCh37 difficult-to-sequence promoters, “bad promoters”, was generated from Ross et al. (2013) supplementary file 13059_2012_3110_MOESM1_ESM.TXT (link checked 08/31/2020). The GRCh37 bad promoter-derived BED file was then remapped to GRCh38 using the NCBI remapping service (https://www.ncbi.nlm.nih.gov/genome/tools/remap).

### Low Complexity

Twenty-two Low Complexity stratifications were created to identify variants in difficult regions due to different types and sizes of low complexity sequence (e.g., homopolymers, STRs, VNTRs, other locally repetitive sequences). To capture the full spectrum of repeats, we used a python script to extract Simple_repeats and Low_complexity repeats form the UCSC RepeatMasker-generated file (http://hgdownload.soe.ucsc.edu/goldenPath/hg38/database/rmsk.txt.gz, date accessed 07/22/2019) and UCSC TRF-generated file (http://hgdownload.soe.ucsc.edu/goldenPath/hg38/database/simpleRepeat.txt.gz, date accessed 07/22/2019). Non-overlapping complement regions for some stratifications are also provided, as “notin” files.

### Other Difficult

We provide nine stratifications for GRCh37 and six stratifications for GRCh38 representing additional difficult regions that do not fall into the other stratification groups. These regions include: (1) the VDJ recombination components on chromosomes 2, 14, and 22; (2) the MHC on chromosome 6; (3) L1Hs greater than 500 base pairs; (4) reference assembly contigs smaller than 500kb; and (5) gaps in the reference assembly with 15kb slop. In addition, we used alignments of GRCh38 to GRCh37 to identify regions that were expanded or collapsed between reference assembly releases. For GRCh37, we provide regions with alignments of either none or more than one GRCh38 contig. We also provide regions where the hs37d5 decoy sequences align to GRCh37 indicating potentially duplicated regions. We describe the identification of these regions while generating the new small variant benchmark in Wagner et al. (2020). We generated files containing the L1H subset of LINEs greater than 500 base pairs starting with the rmsk.txt.gz file from UCSC (https://hgdownload.cse.ucsc.edu/goldenPath/hg19/database/rmsk.txt.gz) and (http://hgdownload.cse.ucsc.edu/goldenPath/hg38/database/rmsk.txt.gz) then identify entries with “L1H” and select those greater than 500 base pairs long.

### Segmental Duplications

Nine Segmental Duplication stratifications were generated to identify whether variants are in segmental duplications or in regions with non-trivial self-chain alignments.

Non-trivial self-chains are regions where one part of the genome aligns to another due to similarity in sequence, e.g., due to genomic duplication events. Segmental Duplications from UCSC (hgdownload.cse.ucsc.edu/goldenPath/hg38/database/genomicSuperDups.txt.gz, link checked 08/31/20) were processed to generate stratifications of all segmental duplications, segmental duplications greater than 10 kb, and regions >10 kb covered by more than 5 segmental duplications with >99% identity (Bailey et al., 2001). For stratifications that represent non-trivial alignments of the genome reference to itself, excluding ALT loci, the UCSC chainSelf (hgdownload.cse.ucsc.edu/goldenPath/hg38/database/chainSelf.txt.gz, link checked 08/31/20) and chainSelfLink (hgdownload.cse.ucsc.edu/goldenPath/hg38/database/chainSelfLink.txt.gz, link checked 08/31/20) were used. Together these files were used to produce stratifications for all chainSelf regions and regions greater than 10 kb. Non-overlapping complement regions for some stratifications are also provided, as “notin” files.

### Mappability

Four Mappability stratifications were created to stratify variant calls based on genomic region short read mappability. Regions with low mappability for different read lengths and error rates were generated using the GEM mappability program (Derrien et al., 2012) and BEDOPS genomic analysis tools (Neph et al., 2012). Two sets of parameters were used representing low and high stringency short read mappability.

Non-overlapping complement regions for some stratifications are also provided, as “notin” files.

### Union

Four Union stratifications were created to identify whether variants are in, or not in, different general types of difficult regions or in any type of difficult region or complex variant. The all difficult stratification regions, is the union of all tandem repeats, all homopolymers >6 bp, all imperfect homopolymers >10 bp, all difficult to map regions, all segmental duplications, GC <25% or >65%, “Bad Promoters”, and “OtherDifficultregions”. Additionally stratifications are provided for the union of all difficult to map regions and all segmental duplications. For all stratifications, a “notin” non-overlapping complement is provided as “easy” regions for stratification.

## Notes

https://doi.org/10.18434/mds2-2336

### Summary of Updates

Updated data availability section to include DOI for the archived dataset.

