## Supplementary material for "precisionFDA Truth Challenge V2: Calling variants from short- and long-reads in difficult-to-map regions": Submission Methods

#### *Supplemental Material - Submission Methods*

##### Table of Contents

*Genetres* 3

- TZMTX - GENeTres AUTh Illumina ABMD 3
- 8H0ZB - GENeTres AUTh Illumina ABD 3
- NWQ6Y - GENeTres AUTh Illumina BBTBD 3
- K33QJ - GENeTres AUTh Illumina BBDBM 4

*Kishwar Shafin (UCSC)* 4

- RU88N - PEPPER-DeepVariant (ONT) 4
- XC97E - PEPPER-DeepVariant (ONT_9to6) 4
- 7NJ5C - PEPPER-DeepVariant (PacBIO) 4
- CN36L - PEPPER-DeepVariant (PacBIO+ONT) 4
- FFFGB - Margin-PEPPER-DeepVariant (PacBIO) 4

*Andrew Carroll (google)* 6

- BSODP - DeepVariant Illumina 6
- B1S5A - DeepVariant PacBio 6
- K4GT3 - DeepVariant Hybrid PacBio-Illumina 6
- S7K7S - DeepTrio Illumina 6
- KPXT2 - DeepTrio PacBio 6
- ES3XW - DeepVariant Illumina (filtered for precision) 6
- J04SL - DeepVariant PacBio (filtered for precision) 6

*Robert Everleigh (Canadian Center for Computational Genomics)* 7

- YJN61 - McGill - (C3G) submission v1 7

*Gen Li (genetalks)* 8

**2OT9Q - Pacbio-bwaalt-deepvariant(Genetalks)** 8
**7RR8Z - Pacbio-bwaalt-dvMHC(Genetalks)** 8
**VES2R - Pacbio-mm2-deepvariant(Genetalks)** 8
**R9CXN - Ilumina-bwa-dvnm(Genetalks)** 9
**M9KLP - Ilumina-bwaalt-dvnm-gatkMHC(Genetalks)** 9
**UYMUW - Ilumina-bwaalt-gatk(Genetalks)** 9
**TG5TE - Illumina-bwaalt-deepvariant(Genetalks)** 9
**HF8CT - Pacbio-mm2dv-asmmhc(Genetalks)** 9
**WGQ43 - Pacbio-mm2SNP-bwaINDEL-asmMHC(Genetalks)** 9
**YGOTK - Illumina-Pacbio-asmMHC v1(Genetalks)** 10
**32LOW - Illumina-Pacbio-asmMHC v2(Genetalks)** 10

*Jordi Morata (did not identify in results table)* 10
**ISKOS -** 10

*Christian Brueffer (did not identify in results table)* 11
**ISKOS - Illumina-Novoalign-VarDict** 11

*Sinem  Demirkaya-Budak (seven bridges)* 12
**4HL0B - Seven Bridges GRAF - Illumina** 12

*Calvin Hung (Wasai Technology)* 13

*Varun Jain (DRAGEN)* 14

*Mohammad Sahraeian (Roche Sequencing Solutions)* 16
**RN-Illumina: This submission uses Illumina reads** 16
**RN-PacBio: This submission uses PacBio reads** 18
**RN-Illumina-PacBio: This submission uses Illumina and PacBio reads** 20
**RN-Illumina-PacBio-ONT: This submission uses Illumina, PacBio, and Oxford Nanopore (ONT) reads** 23

*Chirag Jain (TryHard)* 26
**13678  - Winnowmap-bwamem-deepvariant** 26
**PGXA4 - bwamem-deepvariant** 26
**C6JUX - winnowmap-deepvariant** 26

*Mian Umair Ahsan (Wang Genomics Lab)* 27

*Hanying Feng (Sentieon)* 29
**KXBR8 - Illumina short-read-data-only submission** 29
**EIUT6 - PacBio HIFI only submission** 29

**ASJT6 - PacBio HIFI only submission Model2** **29**
**W919C1 - PacBio HIFI submission** **30**
**YBE9U - Combination of Illumina and PacBio HIFI submission Model2** **30**
**WX8VK - Combination of Illumina, PacBio HIFI, and Oxford Nanopore submission** **31**
**CZA1Y - Combination of Illumina, PacBio HIFI, and Oxford Nanopore submission Model2** **31**
**David Jaspez (Genomics Division at ITER)** **32**
SUBMISSION 1: ITER (v1) – Illumina 32
SUBMISSION 2: ITER (v2) – Illumina filtered 32
SUBMISSION 3: ITER (v3) – Illumina + PacBio + ONT 33
SUBMISSION 4: ITER (v4) – Illumina + PacBio + ONT filtered 33
Methods not provided for submissions HB8P3, 4GKR1, MECML, SEX9X, NFT0L, 23O09, QUE7Q, 0GOOR, BARQS, YUI27, 0O7FL, 61YRJ, LR1FD, MT57N, and JIPSI

### Genetres

I used the full hg38/GRCh38 reference genome distributed by 1000 genomes, derived from NCBI set with HLA and decoy alternative alleles.

#### TZMTX - GENeTres AUTh Illumina ABMD

HG002-A-B_M-D.vcf.gz HG003-A-B_M-D.vcf.gz HG004-A-B_M-D.vcf.gz
1. Mapper: BWA, Version: BWA-0.7.17 (r1188), Parameters: bwa mem -c 250 -M -t -R, Link to code: https://github.com/lh3/bwa, Citation: https://arxiv.org/abs/1303.3997
2. Variant caller: DeepVariant, Version: 0.10.0, Parameters: --model_type=WGS, Link to code: https://github.com/google/deepvariant, Citation: https://www.nature.com/articles/nbt.4235
3. Variant filtering: No variant filtering
4. Trimming with Atropos before mapping yields better results and reduces FPs. MarkDuplicates also produces slightly better results. DeepVariant seems to be affected slightly with the additional steps.

#### 8H0ZB - GENeTres AUTh Illumina ABD

HG002-A-B-D.vcf.gz HG003-A-B-D.vcf.gz HG004-A-B-D.vcf.gz

1. Mapper: BWA, Version: BWA-0.7.17 (r1188), Parameters: bwa mem -c 250 -M -t -R, Link to code: https://github.com/lh3/bwa, Citation: https://arxiv.org/abs/1303.3997
2. Variant caller: DeepVariant, Version: 0.10.0, Parameters: --model_type=WGS, Link to code: https://github.com/google/deepvariant, Citation: https://www.nature.com/articles/nbt.4235
3. Variant filtering: No variant filtering
4. Trimming with Atropos before mapping yields better results and reduces FPs. It was an attempt without MarkDuplicates (control). DeepVariant seems to be affected slightly with the additional steps.

##### NWQ6Y - GENeTres AUTh Illumina BBTBD

HG002-BBT-B-D.vcf.gz HG003-BBT-B-D.vcf.gz HG004-BBT-B-D.vcf.gz
BBT means trimming + contaminant filtering + deduplication with BBTools

1. Mapper: BWA, Version: BWA-0.7.17 (r1188), Parameters: bwa mem -c 250 -M -t -R, Link to code: https://github.com/lh3/bwa, Citation: https://arxiv.org/abs/1303.3997
2. Variant caller: DeepVariant, Version: 0.10.0, Parameters: --model_type=WGS, Link to code: https://github.com/google/deepvariant, Citation: https://www.nature.com/articles/nbt.4235
3. Variant filtering: No variant filtering
4. Trimming and contaminant filtering with BBDuc and deduplicate with BBTools dedupe before mapping yields better results and seems to produce better results in indels (BBMap_38.86 with default parameters). DeepVariant seems to be affected slightly with the additional steps.

##### K33QJ - GENeTres AUTh Illumina BBDBM

HG002-BBD-BM-D.vcf.gz HG003-BBD-BM-D.vcf.gz HG004-BBD-BM-D.vcf.gz
BBT means trimming + contaminant filtering + deduplication with BBTools

1. Mapper: BWA, Version: BWA-0.7.17 (r1188), Parameters: bwa mem -c 250 -M -t -R, Link to code: https://github.com/lh3/bwa, Citation: https://arxiv.org/abs/1303.3997
2. Variant caller: DeepVariant, Version: 0.10.0, Parameters: --model_type=WGS, Link to code: https://github.com/google/deepvariant, Citation: https://www.nature.com/articles/nbt.4235
3. Variant filtering: No variant filtering
4. Trimming with BBDuc before mapping seems to yield better results. It was a control attempt without deduplication with BBtools dedupe (BBMap_38.86 with default parameters). DeepVariant seems to be affected slightly with the additional steps.

#### Kishwar Shafin (UCSC)

Submission names:

##### RU88N - PEPPER-DeepVariant (ONT)
##### XC97E - PEPPER-DeepVariant (ONT_9to6)
##### 7NJ5C - PEPPER-DeepVariant (PacBIO)
##### CN36L - PEPPER-DeepVariant (PacBIO+ONT)
##### FFFGB - Margin-PEPPER-DeepVariant (PacBIO)

Mapper:
ONT: minimap2, version: 2.17-r941
PacBIO: pbmm2, version: 1.3.0--h56fc30b_0
Mapper parameters:
ONT: minimap2 -a -z 600,200 -x map-ont -t <threads> <GRCh38.fasta> <reads.fastq> \
| samtools view -hb -F 0x904 > unsorted.bam

```
samtools sort -@38 -o HG00X_ONT_reads_2_GRCh38.bam unsorted.bam
```

PacBIO: --preset CCS

Variant caller:

PEPPER-DeepVariant (https://github.com/kishwarshafin/pepper)

Parameters and details posted here:
https://github.com/kishwarshafin/pepper/blob/master/docs/PEPPER_variant_calling.md

Variant filtering: None

New innovations:

Variant calling with ONT reads is challenging due to the high error-rate of the reads. The existing implementation of DeepVariant uses a deterministic approach to find candidate variants before genotyping. The allele frequency-based candidate finding method fails to work on noisy long reads like ONT as it picks up too many candidates.

In our collaborative effort to call variants with ONT reads, we paired a recurrent neural network-based candidate finder (PEPPER) with DeepVariant. PEPPER is a haploid assembly polisher redesigned to call variants with DeepVariant. To effectively apply PEPPER-DeepVariant on diploid samples, we use WhatsHap to diplotype the reads before variant calling. The method works in two steps:

Haplotyping:

1. We use PEPPER SNP on an unphased alignment file (BAM) to report SNP sites with high-sensitivity.

2. We use DeepVariant SNP custom trained model that can take PEPPER SNP candidates and filter out false-positive candidates while genotyping each candidate correctly. This yields a precise SNP set.

3. We haplotag the read set with WhatsHap using the precise SNP set from step 2. Given the length of the reads and the accuracy of the SNP set, we get a phased alignment file with long phase-blocks.

Haplotype-aware variant-calling:

1. We use PEPPER HP on each haplotype to get haplotype-specific candidates (SNP + INDELS) and then merge the two candidate sets. The feature encoding of PEPPER HP is done in a haplotype aware manner where the model has information from both haplotypes while finding candidates.

2. Finally, we use a DeepVariant ONT custom trained model that can take PEPPER HP candidates and genotype each of the candidates to provide a highly accurate set of variants. The DeepVariant model used in this step utilizes the phasing information present in the alignment file by generating candidate images that are sorted by haplotags.

Please cite: https://doi.org/10.1038/nbt.4235 (DeepVariant)
https://doi.org/10.1101/085050 (WhatsHap)
https://doi.org/10.1186/s13059-019-1709-0 (haplotype aware diplotyping)

#### Andrew Carroll (google)

##### BSODP - DeepVariant Illumina

##### B1S5A - DeepVariant PacBio

##### K4GT3 - DeepVariant Hybrid PacBio-Illumina

##### S7K7S - DeepTrio Illumina

##### KPXT2 - DeepTrio PacBio

##### ES3XW - DeepVariant Illumina (filtered for precision)

##### J04SL - DeepVariant PacBio (filtered for precision)

Mapper:

BWA MEM v0.7.17. Github: https://github.com/lh3/bwa, https://arxiv.org/abs/1303.3997

Command:

```
bwa mem -t 32 references/grch38_bwa_index/genome.fa \
    HG002.novaseq.pcr-free.35x.R1.fastq.gz \
    HG002.novaseq.pcr-free.35x.R2.fastq.gz \
    -R
@RG\tID:HG002.pcr-free\tPL:ILLUMINA\tPU:NONE\tLB:HG002.pcr-free\tSM
:HG002
```

Reference genome:

GRCh38 no ALT (this is good to capture, as I think more nuanced handling of ALT contigs and supplementing ALT contigs in mappings is one of the findings of the competition).

Variant Caller:

DeepVariant experimental release candidate.

Code for DeepVariant is here: https://github.com/google/deepvariant.

Paper : https://www.nature.com/articles/nbt.4235

The refinements for PrecisionFDA, as well as some additional improvements will be released as DeepVariant v1.0, anticipated in the 1st week of September.

Variant filtering:

In all submissions except the 2 high precision submissions, no post filtering is applied. In the case of the high precision entries only, a filter on the Genotype Quality (GQ) of the variant to add a filter field to calls with GQ < 6 is applied.

Highlight new innovations:

#### Robert Everleigh (Canadian Center for Computational Genomics)

##### YJN61 - McGill - (C3G) submission v1

**PACBIO alignment same parameters for HG002, HG003 and HG004**

reference b38 from
ftp://ftp.ncbi.nlm.nih.gov/genomes/all/GCA/000/001/405/GCA_000001405.15_GRCh38/seqs_for_alignment_pipelines.ucsc_ids/GCA_000001405.15_GRCh38_no_alt_analysis_set.fna.gz

```
module load mugqic/SMRTLink/8.0.0 && \
	pbmm2 align -j 16 --sort --preset CCS -L 0.1 -c 0 \
		--rg
	"@RG\tID:HG002\tPL:PACBIO\tDS:READTYPE=CCS\tSM:HG002_35x_PacBio_14kb-15kb\tPM:SEQUELII" \
		Homo_sapiens.GRCh38_primary.fa \
		HG002_35x_PacBio_14kb-15kb.fastq \
		HG002_35x_PacBio_14kb-15kb_primary.sorted.bam
```

**PACBIO variant calling with clair2 v2.1.0 same parameters for HG002, HG003, and HG004**

```
clair.py callVarBamParallel \
	--chkpnt_fn clair/model/pacbio/model \
	--ref_fn Homo_sapiens.GRCh38_primary.fa \
	--bam_fn HG002_35x_PacBio_14kb-15kb.sorted.bam \
	--sampleName HG002 --output_prefix clair2/HG002_pacbio \
	--minCoverage 4 --tensorflowThreads 4 \
	> commandsHG002old.sh
cat commandsHG002old.sh | parallel -j4
```

**Combine 3 vcfs with bcftools 1.9**

```
bcftools merge -Oz \
	-o allSamples.pacbio_primary.clair2.vcf.gz \
	HG002_35x_PacBio_14kb-15kb_primary.clair2.vcf.gz \
	HG003_35x_PacBio_14kb-15kb_primary.clair2.vcf.gz \
	HG004_35x_PacBio_14kb-15kb_primary.clair2.vcf.gz
```

**Filter out mendelian inconsistency using bcftools plugin +mendelian v10.2**

```
	bcftools +mendelian -c -d -Oz\
		-t HG004,HG003,HG002 \
		-o allSamples.pacbio_primary.clair2.mic.vcf.gz \
		allSamples.pacbio_primary.clair2.vcf.gz
```

**Split back to individual sample vcf bcftools plugin +split v10.2**

```
bcftools +split allSamples.pacbio_primary.clair2.mic.vcf.gz \
    -Oz -o split_pb_mic -i'GT="alt"'
```

#### Gen Li (genetalks)

The raw reads were mapped to GRCh38 without alternative contigs using minimap2(2.17-r974-dirty) with parameters '-H -a -x map-pb'. The output SAM file was converted to BAM, then sorted and indexed using samtools 1.10(using htslib 1.10). For non-difficult and diffcult regions excluded MHC, short variants were called using deepvariant(0.10.0) with parameters '--model_type PACBIO'. For MHC region, an assembly-based approach combined with an in-house haplotype-aware caller were used to do variants calling. HiFi reads mapped to MHC region were extracted and assembled by hifiasm (0.7-dirty-r255). The assembled two haplotypes were remapped to chromosome 6 using minimap2 (2.17-r974-dirty) with parameters '-x asm10'. The candidate events were built from haplotype alignments. Candidate adjacent variants with a distance smaller than 2 were merged into either MNPs or complex variants in terms of haplotype phase information. Also, a right shift operation was performed to simplify the variants representation for complex mixed variants. The right shift algorithm also resolved some local misalignments in low complexity regions. For poorly assembled regions, we recall variants in these regions using above haplotype-aware caller from raw HiFi reads and finally two callsets were merged. The implementation of calling algorithm for MHC region can be found at: https://github.com/Genetalks/callMHC.git.

The calling algorithm innovatively used hifiasm, a versatile tool for diploid assembly based on Pacbio HiFi reads, to assembly MHC region. Comparing to approaches like trio-binning, the method can be applied to single sample without parental sequencing data. Comparing to two-phase assembly approaches, which call confident variants at first and assemble binned reads separately for two parental haplotypes, the method is more efficient with competitive accuracy. Furthermore, the method can also be applied to other regions where reference lacks diversity like MHC.

Submission specific methods

##### 2OT9Q - Pacbio-bwaalt-deepvariant(Genetalks)

1. Mapper: BWA alt
2. Variant Caller: DeepVariant
3. Variant Filtering
4. Additional Comments

##### 7RR8Z - Pacbio-bwaalt-dvMHC(Genetalks)

1. Mapper: BWA alt
2. Variant Caller: DeepVariant, with MHC specific model trained and used for MHC
3. Variant Filtering
4. Additional Comments

#### VES2R - Pacbio-mm2-deepvariant(Genetalks)

1. Mapper: minimap2
2. Variant Caller: DeepVariant
3. Variant Filtering
4. Additional Comments

#### R9CXN - Ilumina-bwa-dvnm(Genetalks)

1. Mapper: bwa, with GATK best practices used for pre-processing
2. Variant Caller: DeepVariant; trained new model for difficult regions
3. Variant Filtering
4. Additional Comments

#### M9KLP - Ilumina-bwaalt-dvnm-gatkMHC(Genetalks)

1. Mapper: BWA alt
2. Variant Caller: DeepVariant using new difficult regions model, GATK used to call variants in MHC
3. Variant Filtering
4. Additional Comments

#### UYMUW - Ilumina-bwaalt-gatk(Genetalks)

Followed GATK best practices

1. Mapper: bwa alt
2. Variant Caller: GATK
3. Variant Filtering
4. Additional Comments

#### TG5TE - Illumina-bwaalt-deepvariant(Genetalks)

1. Mapper: bwa alt
2. Variant Caller: DeepVariant
3. Variant Filtering
4. Additional Comments

#### HF8CT - Pacbio-mm2dv-asmmhc(Genetalks)

1. Mapper: minimap2
2. Variant Caller: DeepVariant, hifiasm assembly used for MHC
3. Variant Filtering
4. Additional Comments

#### WGQ43 - Pacbio-mm2SNP-bwaINDEL-asmMHC(Genetalks)

Haplotype-aware deepvariant

1. Mapper: minimap2
2. Variant Caller: DeepVariant, hifiasm assembly used for MHC
3. Variant Filtering

4. Additional Comments

##### YGOTK - Illumina-Pacbio-asmMHC v1(Genetalks)
Combined best 2 callsets from ILMN and PacBio HiFi

1. Mapper:
2. Variant Caller: DeepVariant
3. Variant Filtering
4. Additional Comments

##### 32LOW - Illumina-Pacbio-asmMHC v2(Genetalks)
Combined best 2 callsets from ILMN and PacBio HiFi

1. Mapper:
2. Variant Caller:
3. Variant Filtering
4. Additional Comments

#### Jordi Morata (did not identify in results table)
##### ISKOS -

1. Mapper, version, parameters, and link to code, and citation if applicable

BWA MEM 0.7.17. Duplicated reads were marked with Picard and base quality score was recalibrated with GATK's BaseQualityScoreRecalibrator.

2. Variant caller, version, parameters, and link to code, and citation if applicable
3. Variant filtering and/or merging, if applicable

Two different softwares were used to call variants on the resulting alignment files (BAMs):

1) Variant calling was performed with GATK (v4.1.7.0) HaplotypeCaller for the three individuals of the trio (HG002, HG003, HG004). Individual gVCFs were then combined into a multisample gVCF with CombineGVCFs. A multisample VCF was generated with GenotypeGVCFs and single sample VCFs were obtained with bcftools. Variants likely to be artifacts were tagged with GATK-VariantFiltration (BaseQRankSum > 4.0, BaseQRankSum < -4.0; FS > 60 ; FS > 200; ReadPosRankSum < -8; ReadPosRankSum > 20; MQRankSum < -12.5 ; QD < 2; MQ < 40). Normalization of variants was performed with bcftools norm, obtaining the VCF file with the alias HC_raw. A subset of those variants with DP>7 was stored in a separate file (HC_DPgt7).

2) Joint Variant calling with all three samples was performed with Strelka 2 (v2.9.9) obtaining a multisample VCF. Single sample VCFs were obtained with bcftools. Normalization of variants was performed with bcftools norm, obtaining the VCF file with

the alias S2_raw. A subset of those variants with DP>7 was stored in a separate file (S2_DPgt7)

We performed the intersection of HaplotypeCaller and Strelka DP filtered results (HC_DPgt7 & S2_DPgt7), selecting those variants which had the status of PASS in at least one of the callers, generating the file Isec_Pass.

The final variants file was obtained reevaluating the variants from HC_raw with the information from Isec_Pass. Variants from HC_raw present in Isec_Pass, regardless of their original filter status, were tagged as PASS. The rest of variants were additionally tagged as 'LowEvidence'.

4. Highlight any new innovations to field that their variant calling method employed (NA)

#### Christian Brueffer (did not identify in results table)

##### H9OJ3 - Illumina-Novoalign-VarDict

The workflow was implemented using Snakemake and conda/bioconda environments.

1. Mapper, version, parameters, and link to code, and citation if applicable

Reads were mapped with Novoalign 4.02.02, and streamed into biobambam bamsormadup for duplicate marking.

Novoalign 4.02.02 (Novocraft, Malaysia)

-c 12 -d GCA_000001405.15_GRCh38_no_alt_analysis_set.fna.ndx -f {fq1} {fq2} -F ILM1.8 -o SAM "@RG\\tID:{id}\\tCN:{center}\\tLB:{sample}\\tPL:ILLUMINA\\tSM:{sample}" --tune NOVASEQ

biobambam 2.0.87 (Tischler and Leonard, Biobambam: Tools for read pair collation based algorithms on BAM files, 2014, Source Code for Biology and Medicine)

bamsormadup inputformat=sam outputformat=bam threads=12 SO=coordinate

2. Variant caller, version, parameters, and link to code, and citation if applicable

VarDict-Java 1.7.0 (Lai et al, VarDict: a novel and versatile variant caller for next-generation sequencing in cancer research, 2016, Nucleic Acids Research)

3. Variant filtering and/or merging, if applicable

Variants were annotated with the data sources dbSNP v151 and gnomAD 2.1, as well as the Danish and Swedish reference genomes using vcfanno.

Variants were filtered using SnpSift to only keep variants in the standard human chromosomes, and then using the SnpSift filter string '(FILTER = 'PASS') & !(TYPE = 'Complex') & ( ((PMEAN >= 6) & (NM <= 1.3) & ((AF >= 0.43 & AF <= 0.57) | (AF >= 0.93))) | ((exists dbSNPb151_ID) | (gnomad_wes_AF >= 0.000001) | (gnomad_wgs_AF >= 0.000001) | (swegen_AF >= 0.0001) | (danish_gen_AF >= 0.0001)) )'.

vcfanno 0.3.2 (Pedersen et al, Vcfanno: fast, flexible annotation of genetic variants, 2016, Genome Biology) SnpSift 4.3.1t (Cingolani et al, Using Drosophila melanogaster as a model for genotoxic chemical mutational studies with a new program, SnpSift, 2012, Frontiers in Genetics)

4. Highlight any new innovations to field that their variant calling method employed

None.

#### Sinem Demirkaya-Budak (seven bridges)

##### 4HL0B - Seven Bridges GRAF - Illumina

You can find the information needed about our submission (Seven Bridges Genomics, Submission ID: 4HL0B) for your manuscript below.

1. Mapper, version, parameters, and link to code, and citation if applicable

GRAF Aligner v1
Command line:

```
aligner
--vcf SBG.Graph.B38.V4.vcf.gz
--reference SBG.Graph.B38.V4.fa
--markdup
--fmt bam
-q HG003.novaseq.pcr-free.35x.R1.fastq.gz
-Q HG003.novaseq.pcr-free.35x.R2.fastq.gz
--read_group_platform 'NovaSeq' --read_group_sample 'HG003'
--sort
--tmp ./sort.chunk.XXXXXX
--sort_mem 30000
--hts_threads 40
--merge_threads 12
--keep  -o HG003.novaseq.pcr-free.35x.R.bam
&& sambamba index -t 36 HG003.novaseq.pcr-free.35x.R.bam
```

For the details of the parameters you can refer to our user guide:
https://hello.sevenbridges.com/hubfs/Graph%20Files/GRAF_Technical_Guide_v1062020.pdf

2. Variant caller, version, parameters, and link to code, and citation if applicable

GRAF Variant Caller v1
Command line:
rasm
-v HG003.novaseq.pcr-free.35x.R.vcf
-a SBG.Graph.B38.V4.bed
-f SBG.Graph.B38.V4.fa
-b HG003.novaseq.pcr-free.35x.R.bam
-g SBG.Graph.B38.V4.vcf.gz
-x all
-s 10
-t 0
For the details of the parameters you can refer to our user guide:
https://hello.sevenbridges.com/hubfs/Graph%20Files/GRAF_Technical_Guide_v1062020.pdf

3. Variant filtering and/or merging, if applicable

None

4. Highlight any new innovations to field that their variant calling method employed

For any mention of our pipeline, you can cite this article: Rakocevic, Goran, et al. "Fast and accurate genomic analyses using genome graphs." Nature genetics 51.2 (2019): 354-362. https://www.nature.com/articles/s41588-018-0316-4

#### Calvin Hung (Wasai Technology)

1. bwa-mem 0.7.17 with Wasai-Lightning™ FPGA accelerator
2. GATK 4.1.3.0 with Wasai-Lightning™ FPGA accelerator
3. N/A
4. Wasai-Lightning™ FPGA accelerator for BWA-MEM + GATK

#### Varun Jain (DRAGEN)

### DRAGEN EXPERIMENTAL EXTENSION INTO DIFFICULT REGIONS – ILLUMINA READS

**DRAGEN team at Illumina:**
Mike Ruehle, Varun Jain, Bryan Lajoie, Cooper Roddey, Severine Catreux, Rami Mehio

**Data type used:** Illumina
**Number of samples required:** Single sample method

**Input preparation protocol:**
1) Map reads with DRAGEN 3.7.x Mapper, run in alt-aware mode
2) Reference used: hg38 (augmented with population haplotypes)
3) De-duplicate reads with DRAGEN duplicate marker
4) DRAGEN small Variant Caller with DRAGEN 3.7.x

**Background:**
DRAGEN is Illumina's ultra-fast and highly accurate secondary analysis platform, which supports a wide array of pipelines, including mapping and aligning, sort, dedup, and multiple variant callers. The small Variant Caller methods are described in DRAGEN Accuracy Application Note (https://science-docs.illumina.com/documents/Informatics/dragen-v3-accuracy-appnote-html-970-2019-006/Content/Source/Informatics/Dragen/dragen-v3-accuracy-appnote-970-2019-006/dragen-v3-accuracy-appnote-970-2019-006.html). The most recently released DRAGEN version is 3.6.3. To open-source its methods and unify secondary analysis pipelines, Illumina and the Broad have a collaboration to co-develop analysis methods and pipelines. As part of this collaboration, several improvements from the DRAGEN small variant caller will be available in the upcoming open-source DRAGEN-GATK release, planned for Q3, 2020 (cf. https://gatk.broadinstitute.org/hc/en-us/articles/360039984151-DRAGEN-GATK-Update-Let-s-get-more-specific).

**Improvements added to this submission:**
This submission incorporates several new features and capabilities that extend DRAGEN's capabilities beyond the 3.6 release. We used the context of this challenge to explore new frontiers in the accuracy achievable with short reads, specifically in the extended regions of the new v4.1 truth set. The improvements have various levels of maturity: some improvements have been under development for a few months and are ripe to be deployed in upcoming DRAGEN releases, while others should be considered experimental proofs-of-concepts, leading us to further exploration and development.

- **Overlapping Variant Detection:** This improvement refers to jointly genotyping potentially overlapping variants at multiple loci in a single region for a single sample. This yields both SNP and INDEL accuracy improvement over genotyping a single locus at a time.

- **Experimental leveraging of population haplotypes for improved mapping accuracy:** In this experimental step, we augmented the hg38 reference with several hundred thousand short alternate contigs derived from population haplotypes, effectively evolving it towards a graph reference, and used the DRAGEN Mapper's alt-awareness capabilities to project reads matching these population haplotypes to corresponding primary assembly alignments. This improved mapping accuracy and variant calls in difficult-to-map regions.

Relative to DRAGEN version 3.6, the combination of these improvements yielded a 48% reduction of SNP errors and a 41% reduction of INDEL errors for HG002.

**DRAGEN 3.7.x Command Lines:**

- Below is the DRAGEN command line to generate the HT

```
dragen
--build-hash-table true
--ht-reference hg38.fa
--ht-alt-liftover /opt/edico/liftover/bwa-kit_hs38DH_liftover.sam
--output-directory /tmp/
--ht-num-threads 40
--ht-pop-alt-contigs /opt/edico/liftover/pop_altcontig.fa
--ht-pop-alt-liftover /opt/edico/liftover/pop_liftover.sam
--ht-pop-snps /opt/edico/liftover/pop_snps.vcf
```

- DRAGEN command line for Fastq □ VCF

```
dragen
-1 HG002.novaseq.pcr-free.35x.R1.fastq.gz
-2 HG002.novaseq.pcr-free.35x.R2.fastq.gz
-f -r pop_HT
--enable-map-align true
--enable-map-align-output true
--enable-sort true
--enable-duplicate-marking true
--enable-variant-caller true
--vc-enable-joint-detection true
--output-directory /tmp/
--output-file-prefix HG002
--RGID DRAGEN_RGID --RGSM DRAGEN
```

A BSSH app to run these DRAGEN command lines is currently being developed.

#### Mohammad Sahraeian (Roche Sequencing Solutions)

#### **RN-Illumina:** This submission uses Illumina reads

1-Mapper:

- Alignment:
  - Tool: BWA-MEM
  - version: 0.7.15
  - Parameters: -M
  - Code: https://sourceforge.net/projects/bio-bwa/files/
  - Citation: Li, H. Aligning sequence reads, clone sequences and assembly contigs with BWA-MEM. https://arxiv.org/abs/1303.3997 (2013).
- FASTQ and BAM process:
  - FASTQ Trimming:
    - Tool: AlienTrimmer
    - Version: 0.4.0
    - Code: ftp://ftp.pasteur.fr/pub/gensoft/projects/AlienTrimmer/
    - Citation: Criscuolo, A. & Brisse, S. AlienTrimmer: a tool to quickly and accurately trim off multiple short contaminant sequences from high-throughput sequencing reads. Genomics 102, 500–506 (2013).
  - MarkDuplicates:
    - Tool: Picard
    - Version: 2.22.4
    - Code: https://broadinstitute.github.io/picard/
  - IndelRealigner:
    - Tool: GATK
    - Version: 3.8.0
    - Code: https://gatk.broadinstitute.org/hc/en-us
    - Citation: DePristo, M. A. et al. A framework for variation discovery and genotyping using next-generation DNA sequencing data. Nature Genet. 43, 491–498 (2011)

2- Variant Caller:

- The predictions are based on the adaptation of NeuSomatic for germline variant calling. Here we used NeuSomatic in ensemble mode (GATK HaplotypeCaller calls were also used as extra feature channels to the network). We also used the assembly-based GATK HaplotypeCaller calls in the post processing step to resolve network calls that are complex, multi-allelic, or for large indels, and report the final ref/alt.
- Version:
  - NeuSomatic: The version used for submission is not public.
  - GATK: 4.1.7.0
  - Picard: 2.22.4
- Code:
  - NeuSomatic: https://github.com/bioinform/neusomatic
  - GATK: https://gatk.broadinstitute.org/hc/en-us
- Parameters:

- NeuSomatic parameters (the version used for submission is not public):
  - Preprocess.py: --scan_maf 0.01 --min_mapq 10 --snp_min_af 0.06 --snp_min_bq 10 --snp_min_ao 3 --ins_min_af 0.03 --del_min_af 0.03 --scan_window_size 10000 --add_extra_features --window_extend 10000 --max_cluster_size 1000 --num_splits 100
  - postprocess.py: --postprocess_max_dist 10
  - call.py: --batch_size 20
  - train.py: --coverage_thr 150 --batch_size 1000 --lr 0.01 --lr_drop_epochs 400 --max_epochs 1000 --boost_none 10
- GATK parameters:
  - HaplotypeCaller: --native-pair-hmm-threads 1

- Citation:
  - NeuSomatic: Sahraeian, S. M. E. et al. Deep convolutional neural networks for accurate somatic mutation detection. Nat. Commun. 10, 1041 (2019).
  - GATK HaplotypeCaller: Poplin, R. et al. Scaling accurate genetic variant discovery to tens of thousands of samples. Preprint at https://www.bioRxiv.org/content/10.1101/201178v2 (2018).

3- Variant filtering:

- The CNN output yields classification scores for variant-type classification (4 classes: None/Insertion/Deletion/SNP), length classification (0 for non-variants, 1 for SNPs and 1-base deletions, 2 and >=3 for INDELs), and genotype (0/0, 0/1, 1/1, 1/2). Based on these predictions, a variant will be reported along with its quality score for each candidate site. We further resolve the complex, multi-allelic, and large indels in the post processing steps. Variants with quality scores larger than 0.7 are reported as PASS calls.

4-Highlights of algorithm:

- We used an extended version of the convolutional neural network (CNN) framework in NeuSomatic for germline variant calling in this submission. NeuSomatic uses a novel summarization of tumor/normal alignment information as a set of input matrices that can be adopted to efficiently train models that learn how to effectively differentiate true somatic mutations from artifacts. For the precisionFDA truth challenge v2, instead of the input channels devoted to the tumor/normal samples, we used a set of input channels for the input sample. As in NeuSomatic, these channels capture the bases' frequencies around a candidate variant, as well as alignment information such as coverage, base quality, mapping quality, strand-bias, and clipping information for reads supporting different bases. We also used separate channels to record information reported by GATK HaplotypeCaller, namely one channel that defines whether GATK HaplotypeCaller has called this variant, and four channels to capture the Genotype Quality (GQ) and the reported genotype of the variant. The same network is used as in NeuSomatic, but with an additional classifier that predicts the genotype (from four classes of 0/0, 0/1, 1/1, and 1/2).
- Steps used for this submission:
  a. AlienTrimmer to trim low-qual and adapter sequences
  b. Reads were aligned using BWA-MEM
  c. Picard MarkDuplicates for marking PCR duplicate read pairs.

d. GATK IndelRealigner to perform indel realignment for improved sequence alignment.
e. Extract GATK HaplotypeCaller's calls (to be used as additional features to input matrices)
f. Scan the indel-realigned BAM, identify candidate variants, and prepare candidate matrices to train the CNN.
g. Train the network on high confidence regions of HG002. For the initial training, the network was trained on chromosomes 1 to 19 and parameters were fine-tuned using evaluations on chromosomes 20 to 22. In the final submission, chromosomes 1 to 22 were used in the training.
h. Infer the germline variants for HG003 and HG004 using the trained network model.
i. Postprocess to report final variants: When the predicted variants were complex, multi-allelic, or large indels, the assembly-based calls from HaplotypeCaller were leveraged to resolve the final set of alleles at the variant locations.

#### **RN-PacBio:** This submission uses PacBio reads

1-Mapper:

- Alignment:
  - Tool: minimap2
  - Version: 2.17
  - Parameters: -a -z 400,50 -k 19 -O 5,56 -E 4,1 -B 5 -r 2k --secondary=no
  - Code: https://github.com/lh3/minimap2
  - Citation: Li, H. (2018). Minimap2: pairwise alignment for nucleotide sequences. Bioinformatics, 34:3094-3100. doi:10.1093/bioinformatics/bty191

2- Variant Caller:

- The predictions are based on the adaptation of NeuSomatic for germline variant calling. Here we used NeuSomatic in ensemble mode (GATK HaplotypeCaller calls were also used as extra feature channels to the network). We also used the assembly-based GATK HaplotypeCaller calls in the post processing step to resolve network calls that are complex, multi-allelic, or for large indels, and report the final ref/alt.
- Version:
  - NeuSomatic: The version used for submission is not public.
  - GATK: 4.1.7.0
- Code:
  - NeuSomatic: https://github.com/bioinform/neusomatic
  - GATK: https://gatk.broadinstitute.org/hc/en-us
- Parameters:
  - NeuSomatic parameters (The version used for submission is not public.):
    - Preprocess.py: --scan_maf 0.01 --min_mapq 1 --min_dp 3 --snp_min_af 0.06 --snp_min_bq 10 --snp_min_ao 3 --ins_min_af 0.06 --del_min_af 0.06 --del_merge_min_af 0.04 --ins_merge_min_af 0.04 --scan_window_size 10000 --add_extra_features --window_extend 10000 --max_cluster_size 1000 --num_splits 100

      - postprocess.py: --postprocess_max_dist 10
      - call.py: --batch_size 20
      - train.py: --coverage_thr 150 --batch_size 1000 --lr 0.01 --lr_drop_epochs 400 --max_epochs 1000 --boost_none 10
    - GATK parameters:
      - HaplotypeCaller: --read-filter MappingQualityReadFilter --read-filter NotSecondaryAlignmentReadFilter --read-filter NotSupplementaryAlignmentReadFilter --pcr-indel-model AGGRESSIVE --native-pair-hmm-threads 1
- Citation:
  - NeuSomatic: Sahraeian, S. M. E. et al. Deep convolutional neural networks for accurate somatic mutation detection. Nat. Commun. 10, 1041 (2019).
  - GATK HaplotypeCaller: Poplin, R. et al. Scaling accurate genetic variant discovery to tens of thousands of samples. Preprint at https://www.bioRxiv.org/content/10.1101/201178v2 (2018).

3- Variant filtering:

- The CNN output yields classification scores for variant-type classification (4 classes: None/Insertion/Deletion/SNP), length classification (0 for non-variants, 1 for SNPs and 1-base deletions, 2 and >=3 for INDELs), and genotype (0/0, 0/1, 1/1, 1/2). Based on these predictions, a variant will be reported along with its quality score for each candidate site. We further resolve the complex, multi-allelic, and large indels in the post processing steps. Variants with quality scores larger than 0.7 are reported as PASS calls.

4-Highlights of algorithm:

- We used an extended version of the convolutional neural network (CNN) framework in NeuSomatic for germline variant calling in this submission. NeuSomatic uses a novel summarization of tumor/normal alignment information as a set of input matrices that can be adopted to efficiently train models that learn how to effectively differentiate true somatic mutations from artifacts. For the precisionFDA truth challenge v2, instead of the input channels devoted to the tumor/normal samples, we used a set of input channels for the input sample. As in NeuSomatic, these channels capture the bases frequencies around a candidate variant, as well as some alignment information such as coverage, base quality, mapping quality, strand-bias, and clipping information for reads supporting different bases. We also used separate channels to record information reported by GATK HaplotypeCaller, namely one channel that defines whether GATK HaplotypeCaller has called this variant, and four channels to capture the Genotype Quality (GQ) and the reported genotype of the variant. The same network is used as in NeuSomatic, but with an additional classifier that predicts the genotype (from four classes of 0/0, 0/1, 1/1, and 1/2).
- Steps used for this submission:
  a. Reads are aligned using minimap2
  b. Extract GATK HaplotypeCaller's calls (to be used as additional features to input matrices)
  c. Scan the input BAM, identify candidate variants, and prepare candidate matrices to train the CNN.

d. Train the network on high confidence regions of HG002. For the initial training, the network was trained using chromosomes 1 to 19 and parameters were fine-tuned using evaluations on chromosomes 20 to 22. In the final submission, chromosomes 1 to 22 were used for training.
e. Infer the germline variants for HG003 and HG004 using the trained network model.
f. Postprocess to report final variants: For the predicted multi-allelic SNVs, the assembly-based calls from HaplotypeCaller were leveraged to resolve the final alleles at variant locations.

##### **RN-Illumina-PacBio:** This submission uses Illumina and PacBio reads

1-Mapper:

- Illumina Alignment:
  - Tool: BWA-MEM
  - version: 0.7.15
  - Parameters: -M
  - Code: https://sourceforge.net/projects/bio-bwa/files/
  - Citation: Li, H. Aligning sequence reads, clone sequences and assembly contigs with BWA-MEM. https://arxiv.org/abs/1303.3997 (2013).
- Illumina FASTQ and BAM process:
  - FASTQ Trimming:
    - Tool: AlienTrimmer
    - Version: 0.4.0
    - Code: ftp://ftp.pasteur.fr/pub/gensoft/projects/AlienTrimmer/
    - Citation: Criscuolo, A. & Brisse, S. AlienTrimmer: a tool to quickly and accurately trim off multiple short contaminant sequences from high-throughput sequencing reads. Genomics 102, 500–506 (2013).
  - MarkDuplicates:
    - Tool: Picard
    - Version: 2.22.4
    - Code: https://broadinstitute.github.io/picard/
  - IndelRealigner:
    - Tool: GATK
    - Version: 3.8.0
    - Code: https://gatk.broadinstitute.org/hc/en-us
    - Citation: DePristo, M. A. et al. A framework for variation discovery and genotyping using next-generation DNA sequencing data. Nature Genet. 43, 491–498 (2011)
- PacBio Alignment:
  - Tool: minimap2
  - Version: 2.17

- Parameters: -a -z 400,50 -k 19 -O 5,56 -E 4,1 -B 5 -r 2k --secondary=no
- Code: https://github.com/lh3/minimap2
- Citation: Li, H. (2018). Minimap2: pairwise alignment for nucleotide sequences. Bioinformatics, 34:3094-3100. doi:10.1093/bioinformatics/bty191

2- Variant Caller:

- The predictions are based on the adaptation of NeuSomatic for germline variant calling (using multiple BAMs). In this submission, separate input channels were used for PacBio and Illumina reads. Here we used NeuSomatic in ensemble mode (GATK HaplotypeCaller calls for Illumina and PacBio reads were also used as extra feature channels to the network). We also used the assembly-based GATK HaplotypeCaller calls in the post processing step to resolve network calls that are complex, multi-allelic, or for large indels, and report the final ref/alt.
- Version:
  - NeuSomatic: The version used for submission is not public.
  - GATK: 4.1.7.0
  - Picard: 2.22.4
- Code:
  - NeuSomatic: https://github.com/bioinform/neusomatic
  - GATK: https://gatk.broadinstitute.org/hc/en-us
- Parameters:
  - NeuSomatic parameters (The version used for submission is not public.):
    - Preprocess.py: --scan_maf 0.01 0.01 --min_mapq 10 1 --min_dp 5 3 --snp_min_af 0.06 0.06 --snp_min_bq 10 --snp_min_ao 3 3 --ins_min_af 0.03 0.06 --del_min_af 0.03 0.06 --del_merge_min_af 0 0.04 --ins_merge_min_af 0 0.04 --scan_window_size 10000 --add_extra_features --window_extend 10000 --max_cluster_size 1000 --num_splits 100
    - postprocess.py: --postprocess_max_dist 10
    - call.py: --batch_size 20 --multi_bam_i 0 1
    - train.py: --coverage_thr 150 --batch_size 1000 --lr 0.01 --lr_drop_epochs 400 --max_epochs 1000 --n_multi_bam 2 --boost_none 10
  - GATK parameters:
    - HaplotypeCaller:

      Illumina: --native-pair-hmm-threads 1
      PacBio: --read-filter MappingQualityReadFilter --read-filter NotSecondaryAlignmentReadFilter --read-filter NotSupplementaryAlignmentReadFilter --pcr-indel-model AGGRESSIVE --native-pair-hmm-threads 1

Citation:
NeuSomatic: Sahraeian, S. M. E. et al. Deep convolutional neural networks for accurate somatic mutation detection. Nat. Commun. 10, 1041 (2019).
GATK HaplotypeCaller: Poplin, R. et al. Scaling accurate genetic variant discovery to tens of thousands of samples. Preprint at https://www.bioRxiv.org/content/10.1101/201178v2 (2018).

3- Variant filtering:

- The CNN output yields classification scores for variant-type classification (4 classes: None/Insertion/Deletion/SNP), length classification (0 for non-variants, 1 for SNPs and 1-base deletions, 2 and >=3 for INDELs), and genotype (0/0, 0/1, 1/1, 1/2). Based on these predictions, a variant will be reported along with its quality score for each candidate site. We further resolve the complex, multi-allelic, and large indels in the post processing steps. Variants with quality scores larger than 0.7 are reported as PASS calls.

4-Highlights of algorithm:

- We used an extended version of the convolutional neural network (CNN) framework in NeuSomatic for germline variant calling (using multiple BAMs) in this submission. NeuSomatic uses a novel summarization of tumor/normal alignment information as a set of input matrices that can be adopted to efficiently train models that learn how to effectively differentiate true somatic mutations from artifacts. For the precisionFDA truth challenge v2, instead of the input channels devoted to the tumor/normal samples, we used a set of input channels for the input sample. When we have multiple alignment BAMs as input (like here where we have Illumina and PacBio BAMs), we devote a set of independent input channels for each BAM. As in NeuSomatic, for each alignment BAM input, these channels capture the bases frequencies around a candidate variant, as well as some alignment information such as coverage, base quality, mapping quality, strand-bias, and clipping information for reads supporting different bases. The channels are aligned across BAM files, such that the bases in each column correspond to the same alignment column in all input BAMs. We also used separate channels to record information reported by GATK HaplotypeCaller for PacBio and Illumina, namely one channel that defines whether GATK HaplotypeCaller has called this variant, and four channels to capture the Genotype Quality (GQ) and the reported genotype of the variant. The same network is used as in NeuSomatic, but with an additional classifier that predicts the genotype (from four classes of 0/0, 0/1, 1/1, and 1/2).
- Steps used for this submission:
    a. Illumina FASTQ processing: AlienTrimmer to trim low-qual and adapter sequences
    b. Illumina Reads are aligned using BWA-MEM
    c. Picard MarkDuplicates on Illumina BAM
    d. GATK IndelRealigner on Illumina BAM
    e. PacBio Reads are aligned using minimap2
    f. Extract GATK HaplotypeCaller's calls for both Illumina and PacBio (to be used as additional features to input matrices)
    g. Scan both the Illumina indel-realigned BAM and PacBio BAM, and prepare candidate matrices to train the CNN. For each candidate location, we have a set of input channels from PacBio and a set of input channels from Illumina, capturing alignment information as in NeuSomatic. GATK Illumina and PacBio calls will also be used as extra feature channels as in NeuSomatic-ensemble mode.
    h. Train the network on high confidence regions of HG002. For the initial training, the network was trained using chromosomes 1 to 19 and parameters were fine-tuned using evaluations on chromosomes 20 to 22. In the final submission, chromosomes 1 to 22 were used for training.

i. Infer the germline variants for HG003 and HG004 using the trained network model.
j. Postprocess to report final variants: For the predicted complex, multi-allelic, and large indels, the Illumina's assembly-based calls from HaplotypeCaller was leveraged to resolve the final ref/alt. For the predicted multi-allelic SNVs, the PacBio assembly-based calls from HaplotypeCaller were leveraged to resolve the final alleles at the variant locations.

##### **RN-Illumina-PacBio-ONT:** This submission uses Illumina, PacBio, and Oxford Nanopore (ONT) reads

1-Mapper:

- Illumina Alignment:
    - Tool: BWA-MEM
    - version: 0.7.15
    - Parameters: -M
    - Code: https://sourceforge.net/projects/bio-bwa/files/
    - Citation: Li, H. Aligning sequence reads, clone sequences and assembly contigs with BWA-MEM. https://arxiv.org/abs/1303.3997 (2013).
- Illumina FASTQ and BAM process:
    - FASTQ Trimming:
        - Tool: AlienTrimmer
        - Version: 0.4.0
        - Code: ftp://ftp.pasteur.fr/pub/gensoft/projects/AlienTrimmer/
        - Citation: Criscuolo, A. & Brisse, S. AlienTrimmer: a tool to quickly and accurately trim off multiple short contaminant sequences from high-throughput sequencing reads. Genomics 102, 500–506 (2013).
    - MarkDuplicates:
        - Tool: Picard
        - Version: 2.22.4
        - Code: https://broadinstitute.github.io/picard/
    - IndelRealigner:
        - Tool: GATK
        - Version: 3.8.0
        - Code: https://gatk.broadinstitute.org/hc/en-us
        - Citation: DePristo, M. A. et al. A framework for variation discovery and genotyping using next-generation DNA sequencing data. Nature Genet. 43, 491–498 (2011)
- PacBio Alignment:
    - Tool: minimap2
    - Version: 2.17

  - Parameters: -a -z 400,50 -k 19 -O 5,56 -E 4,1 -B 5 -r 2k --secondary=no
  - Code: https://github.com/lh3/minimap2
  - Citation: Li, H. (2018). Minimap2: pairwise alignment for nucleotide sequences. Bioinformatics, 34:3094-3100. doi:10.1093/bioinformatics/bty191
- Oxford Nanopore Alignment:
  - Tool: minimap2
  - Version: 2.17
  - Parameters: 2.17, -a -z 600,200 -x map-ont
  - Code: https://github.com/lh3/minimap2
  - Citation: Li, H. (2018). Minimap2: pairwise alignment for nucleotide sequences. Bioinformatics, 34:3094-3100. doi:10.1093/bioinformatics/bty191

2- Variant Caller:

- The predictions are based on the adaptation of NeuSomatic for germline variant calling (using multiple BAMs). In this submission separate input channels were used for PacBio, Illumina, and ONT reads. Here we used NeuSomatic in ensemble mode (GATK HaplotypeCaller calls for Illumina and PacBio reads were also used as extra feature channels to the network). We also used the assembly-based GATK HaplotypeCaller calls in the post processing step to resolve network calls that are complex, multi-allelic, or for large indels, and report the final ref/alt.
- Version:
  - NeuSomatic: The version used for submission is not public.
  - GATK: 4.1.7.0
  - Picard: 2.22.4
- Code:
  - NeuSomatic: https://github.com/bioinform/neusomatic
  - GATK: https://gatk.broadinstitute.org/hc/en-us
- Parameters:
  - NeuSomatic parameters (The version used for submission is not public.):
    - Preprocess.py: --scan_maf 0.01 0.01 0.1 --min_mapq 10 1 10 --min_dp 5 3 5 --snp_min_af 0.06 0.06 0.1 --snp_min_bq 10 --snp_min_ao 3 3 5 --ins_min_af 0.03 0.06 0.2 --del_min_af 0.03 0.06 0.2 --del_merge_min_af 0 0.04 0.1 --ins_merge_min_af 0 0.04 0.1 --scan_window_size 10000 --add_extra_features --window_extend 10000 --max_cluster_size 1000 --num_splits 100
    - postprocess.py: --postprocess_max_dist 10
    - call.py: --batch_size 20 --multi_bam_i 0 1
    - train.py: --coverage_thr 150 --batch_size 1000 --lr 0.01 --lr_drop_epochs 400 --max_epochs 1000 --n_multi_bam 3 --boost_none 10
  - GATK parameters:
    - HaplotypeCaller:

      Illumina: --native-pair-hmm-threads 1
      PacBio: --read-filter MappingQualityReadFilter --read-filter NotSecondaryAlignmentReadFilter --read-filter NotSupplementaryAlignmentReadFilter --pcr-indel-model AGGRESSIVE --native-pair-hmm-threads 1

Citation:
NeuSomatic: Sahraeian, S. M. E. et al. Deep convolutional neural networks for accurate somatic mutation detection. Nat. Commun. 10, 1041 (2019).
GATK HaplotypeCaller: Poplin, R. et al. Scaling accurate genetic variant discovery to tens of thousands of samples. Preprint at https://www.bioRxiv.org/content/10.1101/201178v2 (2018).
3- Variant filtering:

- The CNN output yields classification scores for variant-type classification (4 classes: None/Insertion/Deletion/SNP), length classification (0 for non-variants, 1 for SNPs and 1-base deletions, 2 and >=3 for INDELs), and genotype (0/0, 0/1, 1/1, 1/2). Based on these predictions, a variant will be reported along with its quality score for each candidate site. We further resolve the complex, multi-allelic, and large indels in the post processing steps. Variants with quality scores larger than 0.7 are reported as PASS calls.

4-Highlights of algorithm:

- We used an extended version of the convolutional neural network (CNN) framework in NeuSomatic for germline variant calling (using multiple BAMs) in this submission. NeuSomatic uses a novel summarization of tumor/normal alignment information as a set of input matrices that can be adopted to efficiently train models that learn how to effectively differentiate true somatic mutations from artifacts. For the precisionFDA truth challenge v2, instead of the input channels devoted to the tumor/normal samples, we used a set of input channels for the input sample. When we have multiple alignment BAMs as input (like here where we have Illumina, PacBio, and ONT BAMs), we devote a set of independent input channels for each BAM. As in NeuSomatic, for each alignment BAM input, these channels capture the bases frequencies around a candidate variant, as well as some alignment information such as coverage, base quality, mapping quality, strand-bias, and clipping information for reads supporting different bases. The channels are aligned across BAM files, such that the bases in each column correspond to the same alignment column in all input BAMs. We also used separate channels to record information reported by GATK HaplotypeCaller for PacBio and Illumina, namely one channel that defines whether GATK HaplotypeCaller has called this variant, and four channels to capture the Genotype Quality (GQ) and the reported genotype of the variant. The same network is used as in NeuSomatic, but with an additional classifier that predicts the genotype (from four classes of 0/0, 0/1, 1/1, and 1/2).
- Steps used for this submission:
    a. Illumina FASTQ processing: AlienTrimmer to trim low-qual and adapter sequences
    b. Illumina Reads are aligned using BWA-MEM
    c. Picard MarkDuplicates on Illumina BAM
    d. GATK IndelRealigner on Illumina BAM
    e. PacBio Reads are aligned using minimap2
    f. ONT Reads are aligned using minimap2
    g. Extract GATK HaplotypeCaller's calls for both Illumina and PacBio (to be used as additional features to input matrices)
    h. Scan the Illumina indel-realigned BAM, PacBio BAM, and ONT BAMs, and prepare candidate matrices to train the CNN. For each candidate location, we have a set of input channels from PacBio and a set of input channels from Illumina, and a set of input channels from ONT capturing alignment information as in

NeuSomatic. GATK Illumina and PacBio calls will also be used as extra feature channels as in NeuSomatic-ensemble mode.

i. Train the network on high confidence regions of HG002. For the initial training, the network was trained using chromosomes 1 to 19 and parameters were fine-tuned using evaluations on chromosomes 20 to 22. In the final submission, chromosomes 1 to 22 were used for training.

j. Infer the germline variants for HG003 and HG004 using the trained network model.

k. Postprocess to report final variants: For the predicted complex, multi-allelic, and large indels, Illumina's assembly-based calls from HaplotypeCaller were leveraged to resolve the final alleles at the variant locations. For the predicted multi-allelic SNVs, the PacBio assembly-based calls from HaplotypeCaller were leveraged to resolve the final alleles at the variant locations.

#### Chirag Jain (TryHard)

##### 13678 - Winnowmap-bwamem-deepvariant

Combined submission PGXA4 (Illumina) indels and C6JUX (HiFi) SNPs

##### PGXA4 - bwamem-deepvariant

1. Mapper: bwa mem default parameters
2. Variant Caller: Deepvariant (0.10.0); using default Illumina model
3. Variant Filtering
4. Additional Comments

##### C6JUX - winnowmap-deepvariant

1. Mapper: winnowmap; https://doi.org/10.1093/bioinformatics/btaa435, https://github.com/marbl/Winnowmap
2. Variant Caller: Deepvariant (0.10.0); using default hifi model
3. Variant Filtering
4. Additional Comments

**Text from Submitter**

**Mapper, version, parameters, and link to code, and citation if applicable**

I had used Winnowmap (v1.01) for hifi reads https://doi.org/10.1093/bioinformatics/btaa435 https://github.com/marbl/Winnowmap ; bwa mem for Illumina PCR-free; default parameters. A special feature in Winnowmap compared to other long-read mappers is that it does not remove highly repetitive k-mers from its index. It employs a special technique based on weighted minimizer sampling that lets it avoid repeat masking without taking a significant hit at performance.

**Variant caller, version, parameters, and link to code, and citation if applicable**
Deepvariant (0.10.0); using default models provided for hifi and Illumina respectively.

**Variant filtering and/or merging, if applicable**
I had provided three vcf files, one that used Illumina reads only, another that used hifi only, and third version that merged variants from both hifi and Illumina. In particular, I had noted that SNP calling accuracy was superior in hifi-based variant call set whereas indel-calling accuracy was superior in Illumina. This is due to the nature of errors in these technologies. Therefore, I pulled SNP calls from the hifi vcf and indel calls from the Illumina vcf to produce the merged vcf.

**Highlight any new innovations to field that their variant calling method employed**
Already highlighted above. (This was mostly me trying to learn more about variant calling tools and benchmarking methods)

#### Mian Umair Ahsan (Wang Genomics Lab)

### Mapper, version, parameters, and link to code, and citation if applicable

We used minimap2 v2.17 for aligning PacBio and ONT reads to GRCh38.

For ONT reads, we used the following minimap2 options:

```
-a -z 600,200 -x map-ont
```

For PacBio reads, we used the following options:

```
-a -x map-pb \
    -k 19 \
    -O 5,56 \
    -E 4,1 \
    -B 5 \
    -z 400,50 \
    -r 2k \
    --eqx \
    --secondary=no
```

#### Variant caller, version, parameters, and link to code, and citation if applicable

We used an ensemble of three variant callers:

1. NanoCaller developed by our lab
   Code: https://github.com/WGLab/NanoCaller (specifically we used v.0.1.0 available at https://github.com/WGLab/NanoCaller/releases/tag/v0.1.0)
   bioarxiv link: https://www.biorxiv.org/content/10.1101/2019.12.29.890418v3

doi: https://doi.org/10.1101/2019.12.29.890418

2. Clair
   Code: https://github.com/HKU-BAL/Clair
   Paper: Luo, R., Wong, C., Wong, Y. *et al.* Exploring the limit of using a deep neural network on pileup data for germline variant calling. *Nat Mach Intell* **2,** 220–227 (2020)
   doi: https://doi.org/10.1038/s42256-020-0167-4

3. medaka
   Code: https://nanoporetech.github.io/medaka/index.html

#### Variant filtering and/or merging, if applicable

We used majority voting between three variant callers to determine variant calls. Quality scores from QUAL field from each variant caller were min-max standardized, and for a given variant call, we assigned it a quality score by taking sum of quality of scores of corresponding variant calls from each variant caller, and dividing it by 3 (if a variant caller did not make any call at a certain genomic position we gave it a quality score of 0).

#### Highlight any new innovations to field that their variant calling method employed

Our internal testing on v4.2 benchmark using RTG vcfeval showed that NanoCaller performs better than medaka and Clair for predicting SNPs on MHC and difficult to map regions, and performs slightly worse than the ensemble. Whereas for indel calling, NanoCaller does not perform as well as other variant callers.

We believe that NanoCaller is able to achieve such high performance for SNP calling because we introduced a novel method of SNP prediction which exploits the long length of the reads and incorporates haplotype information in a deep learning model to predict SNPs. Typically, variant callers such as DeepVariant and Clair, use local information from pileups of bases immediately around a candidate site to determine a variant call. In NanoCaller, we ignore this information, and instead look at the pileups of other SNP sites which we consider highly likely to be heterozygous, that share the same long reads as a given candidate site. The basic idea is this: if a candidate site is a true variant, then if we group the reads at the candidate site according to which base they support, we should see that such grouping of reads is consistent with the haplotype structure of other SNP sites that share the same reads. This allows us to ignore false alleles at a given candidate site if the reads that support the false allele do not consistently support same set of alleles at other SNP sites. Please refer to our bioarxiv paper for more details.

### Hanying Feng (Sentieon)

#### KXBR8 - Illumina short-read-data-only submission

***(A)***

For "Illumina short-read-data-only submission (ID: KXBR8)", we used Sentieon's standard DNAscope pipeline:

1. Mapper, version, parameters, and link to code, and citation if applicable

| Mapper | Version | Parameters | Link to code |
|---|---|---|---|
| Sentieon BWA | 201911 | "-M -K 10000000" | free trial request at www.sentieon.com |

2. Variant caller, version, parameters, and link to code, and citation if applicable

| Variant caller | Version | Parameters | Link to code |
|---|---|---|---|
| Sentieon DNAscope | 201911 | --dbsnp $DBSNP -pcr_indel_model none | free trial request at www.sentieon.com |

3. Variant filtering and/or merging, if applicable
   Sentieon DNAscope

4. Highlight any new innovations to field that their variant calling method employed
   No new development for this challenge in Sentieon DNAscope. We used the standard DNAscope pipeline developed in 2019 based on NIST Truth V3.3.2, but added HG002 Truth V4.1 in the model training. Sentieon DNAscope uses classical statistical methods to identify variant candidates, and machine-learning-filtering model to filter and classify variants.

#### EIUT6 - PacBio HIFI only submission

#### ASJT6 - PacBio HIFI only submission Model2

***(B)***

For "PacBio HIFI only submission (ID: EIUT6)" and "PacBio HIFI only submission Model2 (ID: ASJT6)":

1. Mapper, version, parameters, and link to code, and citation if applicable

| Mapper | Version | Parameters | Link to code |
|---|---|---|---|
| pbmm2 | 1.3.0 | "align -L 0.1 -c 0 --preset CCS --sort" | https://github.com/PacificBiosciences/pbmm2 |

2. Variant caller, version, parameters, and link to code, and citation if applicable

| Variant caller | Version | Parameters | Link to code |
|---|---|---|---|
| Sentieon DNAscope – PacBio HiFi | To be released | To be released in next product upgrade | Currently in engineering mode, to be released in next product upgrade |

3. Variant filtering and/or merging, if applicable
   Sentieon DNAscope

4. Highlight any new innovations to field that their variant calling method employed
   For this challenge, we developed modifications to Sentieon DNAscope to accommodate PacBio HIFI reads. Due to PacBio HIFI's much longer read length, we updated the local assembly and pair-hmm algorithm in DNAscope to examine larger region, hence taking full account of local environment and indel noise characteristics in HIFI reads.
   The difference between the two submissions is the result of training data selection in the machine-learning module for regions outside Truth-V4.1 high-confidence bed file.

#### W919C1 - PacBio HIFI submission

#### YBE9U - Combination of Illumina and PacBio HIFI submission Model2

For "Combination of Illumina and PacBio HIFI submission (ID: W91C1)" and "Combination of Illumina and PacBio HIFI submission Model2 (ID: YBE9U)":

1. Mapper, version, parameters, and link to code, and citation if applicable

| Mapper | Version | Parameters | Link to code |
|---|---|---|---|
| Sentieon BWA | 201911 | "-M -K 10000000" | free trial request at www.sentieon.com |
| pbmm2 | 1.3.0 | "align -L 0.1 -c 0 --preset CCS --sort" | https://github.com/PacificBiosciences/pbmm2 |

2. Variant caller, version, parameters, and link to code, and citation if applicable

| Variant caller | Version | Parameters | Link to code |
|---|---|---|---|
| Sentieon DNAscope | 201911 | --dbsnp $DBSNP -pcr_indel_model none | free trial request at www.sentieon.com |
| Sentieon DNAscope – PacBio HiFi | To be released | To be released in next product upgrade | Currently in engineering mode, to be released in next product upgrade |

3. Variant filtering and/or merging, if applicable
   Sentieon DNAscope

4. Highlight any new innovations to field that their variant calling method employed
   In this challenge, we developed a two-step process to combine the best of two technologies (Illumina and PacBio HiFi). First, we used Sentieon DNAscope to produce variants for Illumina short reads and PacBio HIFI long reads, respectively. This step produced the feature representation of all potential variants from each technology. Second, a machine-learning model was developed and trained based on the concatenated feature vectors from both technologies. The trained model effectively overcomes the limitation of each technology, e.g. low-complexity regions for Illumina short-reads and

homopolymer/tandem-repeat regions for PacBio HIFI. The final result drastically reduced the number of total errors from either individual technology.
The difference between the two submissions is the result of training data selection in the machine-learning module for regions outside Truth-V4.1 high-confidence bed file.

#### WX8VK - Combination of Illumina, PacBio HIFI, and Oxford Nanopore submission

#### CZA1Y - Combination of Illumina, PacBio HIFI, and Oxford Nanopore submission Model2

***(D)***

For regions outside MHC in "Combination of Illumina, PacBio HIFI, and Oxford Nanopore submission (ID:WX8VK)", we used the results from "Combination of Illumina and PacBio HIFI submission (ID: W91C1)" (info provided above)
For regions outside MHC in "Combination of Illumina, PacBio HIFI, and Oxford Nanopore submission Model2 (ID: CZA1Y)", we used the results from "Combination of Illumina and PacBio HIFI submission Model2 (ID: YBE9U)" (info provided above)

For the MHC region in "Combination of Illumina, PacBio HIFI, and Oxford Nanopore submission (ID:WX8VK)" and "Combination of Illumina, PacBio HIFI, and Oxford Nanopore submission Model2 (ID: CZA1Y)":

1. Mapper, version, parameters, and link to code, and citation if applicable

| Mapper | Version | Parameters | Link to code |
|---|---|---|---|
| Sentieon BWA | 201911 | "-M -K 10000000" | free trial request at www.sentieon.com |
| pbmm2 | 1.3.0 | "align -L 0.1 -c 0 --preset CCS --sort" | https://github.com/PacificBiosciences/pbmm2 |
| minimap2 | 2.16 | "-a -z 600,200 -x map-ont" | https://github.com/lh3/minimap2 |

2. Variant caller, version, parameters, and link to code, and citation if applicable

| Variant caller | Version | Parameters | Link to code |
|---|---|---|---|
| Sentieon DNAscope | 201911 | --dbsnp $DBSNP -pcr_indel_model none | free trial request at www.sentieon.com |
| Sentieon DNAscope – PacBio HiFi | To be released | To be released in next product upgrade | Currently in engineering mode, to be released in next product upgrade |
| MAFFT | 7.407 | "--referenceSequenceID 6 --max_gap_length 1000000 --doProduce_pseudoSample 1 | https://mafft.cbrc.jp/alignment/software/ |

| | | |
|---|---|---|
| | | --doProduce_separated VCF 0" |

3. Variant filtering and/or merging, if applicable
   Sentieon DNAscope

4. Highlight any new innovations to field that their variant calling method employed
   We developed a machine learning approach to combine the three call results from the three technologies: 1. we applied Sentieon DNAscope on the Illumina reads; 2. We applied Sentieon DNAscope – PacBio HiFi on the PacBio HiFi reads; 3. We employed Oxford Nanopore reads and Illumina reads to phase the variant call results from PacBio HiFi reads, partitioned the HiFi reads into two phases using the phased variants, then assembled the HiFi reads in each partition, and finally called the variants with the phased assembled contigs using MAFFT. With these three call sets, we applied a machine learning approach to determine the final VCF output.

#### Submissions by the Genomics Division at ITER

##### SUBMISSION 1: ITER (v1) – Illumina

For this submission, we used our in-house variant calling pipeline for Illumina dataset following the Best Practices workflow recommendations for germline variant calling in GATK4. In short, BWA-MEM v0.7.15-r1140 was used to align reads using the GRCh38/hg38 reference genome downloaded from the GATK resource bundle. Then, duplicated reads were marked with the MarkDuplicates tool from GATK. Reads were then processed by a base recalibration tool (BaseRecalibrator tool from GATK) to get a recalibrated BAM. Single nucleotide variants (SNVs) and small insertion and deletion (INDELs) were identified with the HaplotypeCaller (GATK v4.1.4.0) from these BAM files. Finally, SNV and INDEL recalibration was carried out with VariantRecalibrator tool (GATK). We supply a link to our WDL-based pipeline in GitHub: https://github.com/genomicsITER-developers/wdl/tree/master/WGSGermlineSNPsIndels.

##### SUBMISSION 2: ITER (v2) – Illumina filtered

For this submission, we used our in-house variant calling pipeline for Illumina dataset following the Best Practices workflow recommendations for germline variant calling in GATK4. In short, BWA-MEM v0.7.15-r1140 was used to align reads using the GRCh38/hg38 reference genome downloaded from the GATK resource bundle. Then, duplicated reads were marked with the MarkDuplicates tool from GATK. Reads were then processed by a base recalibration tool (BaseRecalibrator tool from GATK) to get a recalibrated BAM. Single nucleotide variants (SNVs) and small insertion and deletion (INDELs) were identified with the HaplotypeCaller (GATK v4.1.4.0) from these BAM files. After that, SNV and INDEL recalibration was carried out with VariantRecalibrator tool (GATK). Finally, we cleaned the VCF file with several filters, such as PASS, QD (Quality by Depth) and MQ (Mapping Quality). We supply a link to our WDL-based pipeline in GitHub: https://github.com/genomicsITER-developers/wdl/tree/master/WGSGermlineSNPsIndels.

#### SUBMISSION 3: ITER (v3) – Illumina + PacBio + ONT

For this submission, we used our in-house variant calling pipeline for Illumina dataset following the Best Practices workflow recommendations for germline variant calling in GATK4. In short, BWA-MEM v0.7.15-r1140 was used to align reads using the GRCh38/hg38 reference genome downloaded from the GATK resource bundle. Then, duplicated reads were marked with the MarkDuplicates tool from GATK. Reads were then processed by a base recalibration tool (BaseRecalibrator tool from GATK) to get a recalibrated BAM. Single nucleotide variants (SNVs) and small insertion and deletion (INDELs) were identified with the HaplotypeCaller (GATK v4.1.4.0) from these BAM files. Finally, SNV and INDEL recalibration was carried out with VariantRecalibrator tool (GATK). We supply a link to our WDL-based pipeline in GitHub: https://github.com/genomicsITER-developers/wdl/tree/master/WGSGermlineSNPsIndels.

For both PacBio and ONT datasets, we built an *ad hoc* pipeline for these technologies using NanoPlot for quality control and Filtlong (v0.2.0) for filtering the reads. Then, minimap2 (v2.17) was used for align reads and Longshot (v0.4.1) for variant calling. After that, we created a consensus VCF file by merging data from the three technologies using the CombineVariants tool from GATK.

#### SUBMISSION 4: ITER (v4) – Illumina + PacBio + ONT filtered

For this challenge, we used our in-house variant calling pipeline for Illumina dataset following the Best Practices workflow recommendations for germline variant calling in GATK4. In short, BWA-MEM v0.7.15-r1140 was used to align reads using the GRCh38/hg38 reference genome downloaded from the GATK resource bundle. Then, duplicated reads were marked with the MarkDuplicates tool from GATK. Reads were then processed by a base recalibration tool (BaseRecalibrator tool from GATK) to get a recalibrated BAM. Single nucleotide variants (SNVs) and small insertion and deletion (INDELs) were identified with the HaplotypeCaller (GATK v4.1.4.0) from these BAM files. Finally, SNV and indel recalibration was carried out with VariantRecalibrator tool (GATK). We supply a link to our WDL-based pipeline in GitHub: https://github.com/genomicsITER-developers/wdl/tree/master/WGSGermlineSNPsIndels.

For both PacBio and ONT datasets, we built an *ad hoc* pipeline for these technologies using NanoPlot for quality control and Filtlong (v0.2.0) for filtering of reads. Then, minimap2 (v2.17) was used for align reads and Longshot (v0.4.1) for variant calling. After that, we cleaned the VCF file with several filters, such as PASS, QD (Quality by Depth) and MQ (Mapping Quality) for Illumina data, and PASS and QUAL for PacBio and ONT data; and created a consensus VCF file by merging data from the three technologies using CombineVariants tool from GATK.
